# *Har-P, a* short *P*-element variant, weaponizes *P*-transposase to severely impair *Drosophila* development

**DOI:** 10.1101/700211

**Authors:** Satyam P. Srivastav, Reazur Rahman, Qicheng Ma, Nelson C. Lau

## Abstract

Without transposon-silencing Piwi-interacting RNAs (piRNAs), transposition causes an ovarian atrophy syndrome in *Drosophila* called gonadal dysgenesis (GD). *Harwich* (*Har*) strains with *P*-elements cause severe GD in F1 daughters when *Har* fathers mate with mothers lacking *P*-element-piRNAs (i.e. *ISO1* strain). To address the mystery of why *Har* induces severe GD, we bred hybrid *Drosophila* with *Har* genomic fragments into the *ISO1* background to create *HISR-D or HISR-N* lines that still cause <u>D</u>ysgenesis or are <u>N</u>on-dysgenic, respectively. In these lines, we discovered a highly truncated *P*-element variant we named “*Har-P*” as the most frequent *de novo* insertion. Although *HISR-D* lines still contain full-length *P*-elements, *HISR-N* lines lost functional *P*-transposase but retained *Har-P*’s that when crossed back to *P*-transposase restores GD induction. Finally, we uncovered *P*-element-piRNA-directed repression on *Har-P’s* transmitted paternally to suppress somatic transposition. The *Drosophila* short *Har-P’s* and full-length *P*-elements relationship parallels the MITEs/DNA-transposase in plants and SINEs/LINEs in mammals.

## INTRODUCTION

The sterility syndrome of “P-M” hybrid dysgenesis in *Drosophila melanogaster* (Engels and Preston, 1979; Kidwell et al., 1977) is due to uncontrolled *P*-element transposition that damages ovarian development and induces female sterility ((Bingham et al., 1982) and reviewed in (Kelleher, 2016)). This gonadal dysgenesis (GD) phenotype occurs in hybrid F1 daughters whose paternal genome comes from a father possessing active *P*-elements (a “*P*” strain) and a maternal genome unable to express *P*-element piRNAs (an “*M*” strain) (Brennecke et al., 2008; Khurana et al., 2011). The fascinating nature of this genetic syndrome is complete fertility in daughters from the reciprocal cross because the mother possessing active *P*-elements contribute *P*-element-derived Piwi-interacting RNAs (piRNAs) to silence these transposons in daughters (Brennecke et al., 2008; Khurana et al., 2011). Thus, despite identical genetic makeup in daughters between the reciprocal crosses, the epigenetic maternal transmission of transposon repression by piRNAs starkly defines female fertility (illustrated in Fig. 1A).

**Figure 1.**
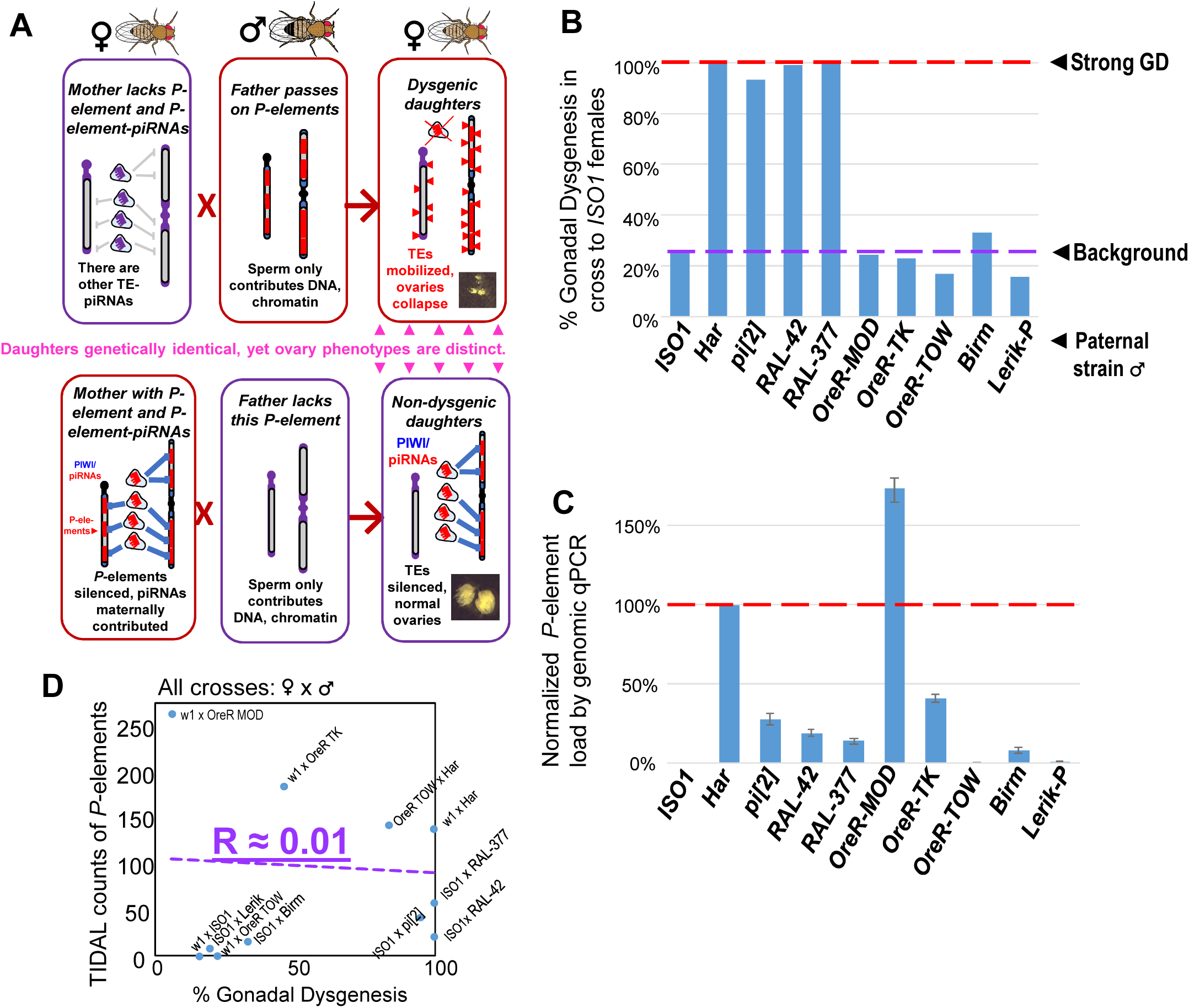
No correlation between paternally-induced Gonadal Dysgenesis (GD) rate and *P*-element copy number. (A) Illustration of the *P*-element-induced GD phenomenon, where two different types of crosses with one parent lacking *P*-elements while the other parents containing *P*-elements can result in genetically identical daughters having very different gonadal phenotypes. (B) GD rates from paternal genome strains mated with *ISO1* females; at least 100 F1 daughters per cross were assayed. (C) Genomic quantitative PCR assessment of *P*-element load of strains, normalized to *Har* at 100%. (D) Scatterplot comparing TIDAL counts of *P*-element insertions to the GD rate reflects the lack of correlation.

Between fertility and complete sterility lies a spectrum of GD induction variation amongst different strain crosses that may be attributed to differential *P*-element copy numbers in different strain genomes (Anxolabehere et al., 1985; Bergman et al., 2017; Biemont et al., 1990; Bingham et al., 1982; Boussy et al., 1988; Kidwell et al., 1981; Ronsseray et al., 1989; Srivastav and Kelleher, 2017; Yoshitake et al., 2018), and capacity to generate piRNAs (Wakisaka et al., 2017). In addition, there are many non-autonomous *P*-element variants that can be mobilized by *P*-transposases, including very short elements from the *pi*[*2*] strain (Bingham et al., 1982; O’Hare and Rubin, 1983) that actually assemble *in vitro* with the *P*-transposase tetramer complex >100X more efficiently than the full-length *P*-element (Tang et al., 2007). However, many earlier studies perceived truncated variants such as the “*KP2*” variants as inhibitors of transposition by acting to titrate *P*-transposase since *P-*element piRNAs were unknown at the time (Black et al., 1987; Gloor et al., 1993; Jackson et al., 1988; Robertson and Engels, 1989; Simmons et al., 2002a). Most studies of GD were typically calibrated with a strong paternal inducer *“P”-*strain like *Harwich* (*Har*) or *pi*[*2*] when mated with “*M*” strain females lacking *P*-elements (Bingham et al., 1982; Brennecke et al., 2008; Kidwell et al., 1977; Rubin et al., 1982). Despite over 40 years of study, what defines a strong paternal inducer of GD has remained a mystery.

Although *P*-element copy numbers in *Har* are significant (120-140 copies, (Khurana et al., 2011)), strains with even more copies like *OreR-MOD* do not induce GD whereas other strong inducer strains that have >75% fewer *P*-element copies than *Har* can also trigger complete GD (Fig. 1B, 1C). Thus, there is a lack of correlation between *P*-element copy number and GD induction (Fig. 1D) that we and others have previously observed (Bergman et al., 2017; Ronsseray et al., 1989; Srivastav and Kelleher, 2017). Since *P*-element copy numbers do not explain GD severity, we hypothesized that a special *P*-element variant or insertion locus might underlie the strong GD phenotype in certain strong “*P*” strains like *Har*. To discover this *P*-element variant, we undertook a reductionist approach to find specific *P*-element variant(s) required for GD induction that revealed unexpectedly a short variant from the *Har* strain that may act together with the full-length *P*-transposase to drive strong GD.

## RESULTS

### GD retention and loss in hybrid Drosophila lines with reduced P-element copy numbers

To genetically isolate the causative transposon strongly inducing GD and facilitate discovery by whole genome sequencing (WGS), we generated hybrid lines where only a minor fraction of the *Har* genome is within the background of the *ISO1* reference genome sequence. We first conducted several fertility-permissive backcrosses between female *Har* and male *ISO1*, selecting hybrid progeny that propagated a red-eye phenotype which we attributed to the “red” eyes due to *Har* alleles replacing the *cn, bw, sp*, alleles on Chromosome 2R (Chr2R) of *ISO1* (Fig. 2A-abridged scheme, Fig. 2-S1-detailed scheme). We then performed an initial GD validation screen with many vials of individual hybrid males crossed to *ISO1* females and selecting for lines that caused 100% GD from this cross. Lines were propagated with additional self-crosses and further in-bred with single-sibling pairs. We then subjected multiple independent *Har-ISO1-Selfed-Red* (*HISR*) lines to a second GD assay. Finally, we conducted qPCR to identify the lines with the greatest reduction of *P*-element copy numbers (Fig. 2B) and settled on 4 lines each either retaining severe paternally-induced GD (*HISR-D*) or had lost this capacity (*HISR-N*) (Fig. 2C).

**Figure 2.**
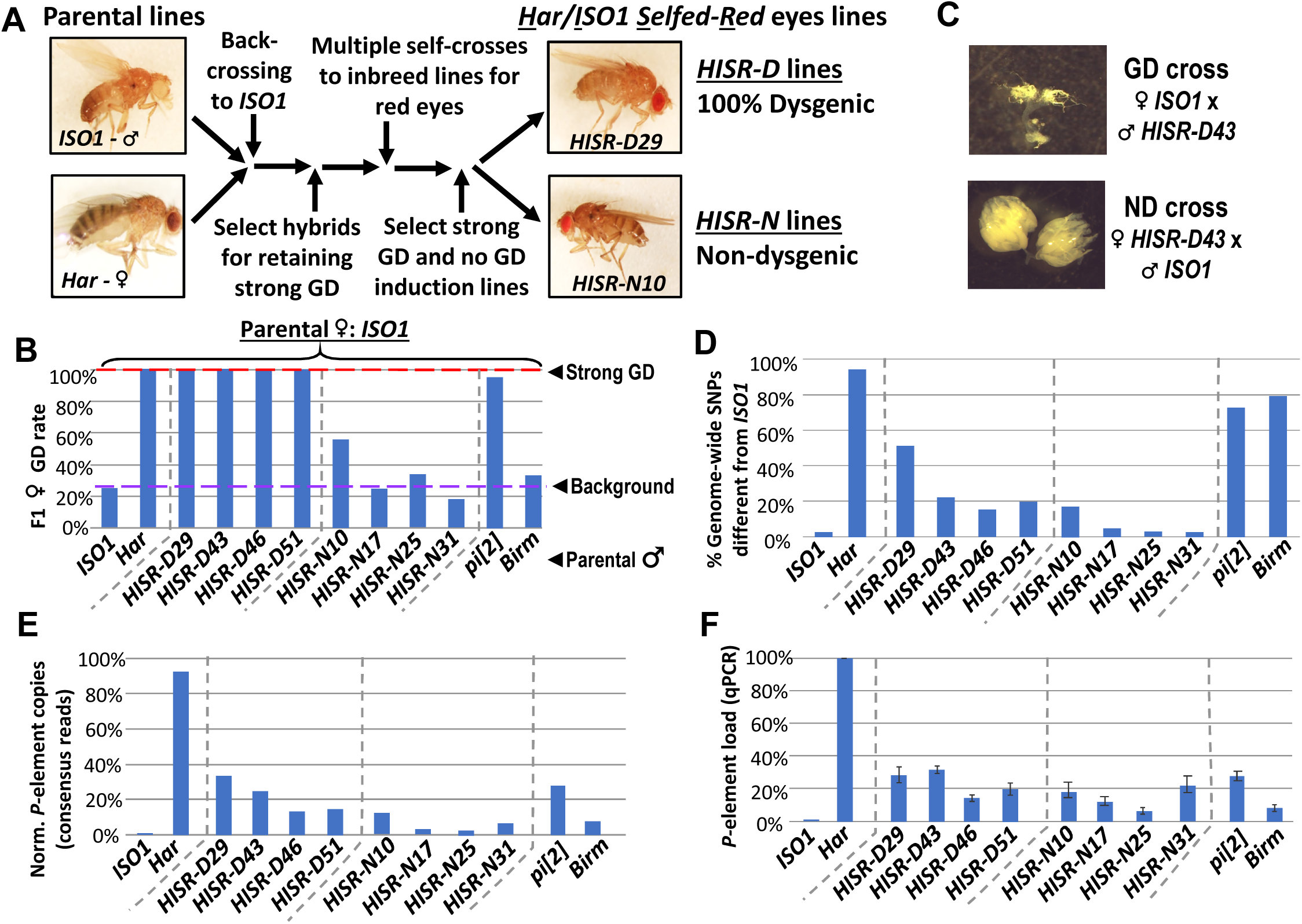
*HISR* lines retaining or losing strong gonadal dysgenesis (GD) induction. (A) Abridged scheme for generating hybrid lines retaining a small fraction of the *Har* genome in the *ISO1* background. Full scheme is Fig. 2 S1. (B) GD rates from paternal genome lines and strains mated with *ISO1* females. (C) Ovarian atrophy phenotype only observed from paternal induction of GD. (D) Genome-wide single nucleotide polymorphism profile differences distinct from *ISO1* genome. (E) Normalized counts of *P*-element copies by consensus read mapping of genomic libraries. (F) qPCR assessment of total genomic *P*-element load.

Genomic PCR genotyping of deletion loci of *Har* compared to *ISO1* in *HISR* lines indicated that these lines carried mostly *ISO1* genomes (Fig. 2-S2). Therefore, we performed WGS of the parental *Har* and *ISO1* strains, the 8 *HISR* lines, and the *pi*[*2*] and *Birmingham* (*Birm*) strains, two classic strains with similar numbers of *P*-elements but diametric capacity to induce GD (Engels et al., 1987; Simmons et al., 1987). Single-Nucleotide Polymorphism (SNP) profiles of *HISR* lines confirmed that only a small percentage of the *Har* genome was retained in mostly an *ISO1* background (Fig. 2D). Quantification of *P*-element copies from WGS with the TIDAL program (Transposon Insertion and Depletion AnaLyzer, Fig.2E) (Rahman et al., 2015) was also consistent with qPCR measurements (Fig. 2F).

### HISR-D lines produce similar levels of P-element piRNAs as the parental Har strain

To determine how substantial reduction in *P*-element copy numbers in *HISR* lines affected *P*-element-directed piRNA production, we generated and sequenced highly-consistent ovarian small RNA libraries (Fig. 3A) and confirmed the expected presence and absence of *P*-element piRNAs in *Har* and *ISO1* ovaries, respectively (Fig. 3B). Surprisingly, there were similar-to-increased levels of *P*-element piRNAs between *HISR-D* and *Har* strains, whereas amongst the *HISR-N* lines, only *HISR-N10* retained *P*-element piRNAs (Fig. 3C). Our own mapping analysis indicated a common 3’ end antisense bias of *P*-element piRNAs that we also confirmed with an independent piRNA analysis pipeline (Han et al., 2015). These mapping patterns are consistent with piRNAs silencing transposons and suppressing hybrid GD (Brennecke et al., 2008; Erwin et al., 2015; Khurana et al., 2011) as well as correlating with all the *HISR-D’s* and *HISR-N10’s* immunity to strong GD induction when these females are mated to *Har* males (Fig. 3D).

**Figure 3.**
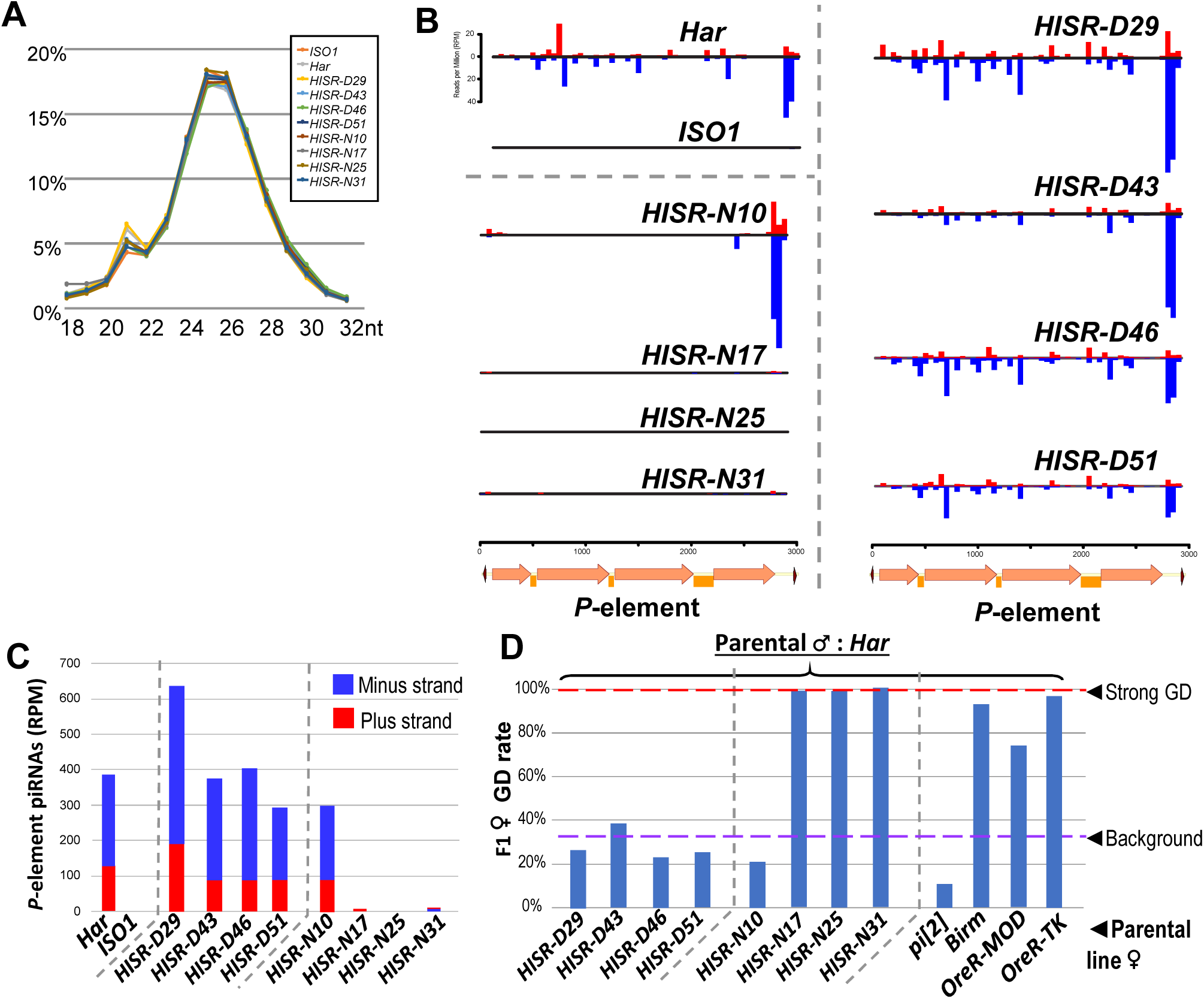
*P*-element directed piRNA production in *HISR* strains ovaries. (A) Nearly identical size distributions of small RNAs from parental and *HISR* ovaries. (B) *P*-element piRNAs coverage plots and (C) quantitation of the *P-*element piRNAs mapping to plus and minus strands, in reads per million (RPM). (D) Assays for repression of *P*-element induced GD for *HISR* strains (N ≥ 100 females) are a good proxy for production of piRNAs silencing *P-*elements.

Additional piRNAs broadly cover the full length of *P*-element in *Har* and *HISR-D* lines (Fig. 3B-top and right), but the notable depletion of internal *P*-element piRNAs in *HISR-N10* (Fig. 3B-middle left) prompted us to conjecture which of its 23 TIDAL-mapped *P*-elements might be stimulating this novel piRNA pattern. We only found one euchromatic *P*-element insertion in *HISR-N10* that specifically coincided with an increase of local piRNAs (Fig. 3-S1A). This *P*-element inserted into the 5’ UTR of *DIP1*, adjacent to the enhancer and promoter region of *Flamenco*, the major piRNA cluster located in a pericentromeric region of the X-chromosome (Brennecke et al., 2007). However, when we selected just the *HISR-N10* X-chromosome balanced with the *FM7a* balancer chromosome (Fig. 3-S1B), this X-chromosome locus did not generate enough *P*-element piRNAs to provide full GD immunity. It is possible for additional *P*-elements to have inserted into major piRNA cluster loci like *42AB*, *Flamenco* and *TAS*-regions as part of the endogenizing process (Khurana et al., 2011; Moon et al., 2018), but the intractable repetitiveness of piRNA cluster regions prevents bioinformatic programs from pinpointing *P*-element insertions in these regions. However, the *P*-element piRNA patterns in *HISR-N10* can be explained by the abundant *P*-element variant that will be discussed below.

### Dispersed P-element landscapes indicate de novo transposition in HISR lines

The selection for “red” eyes of *Har* alleles in *HISR* lines should have replaced the *cn, bw, sp*, alleles on Chromosome 2R (Chr2R) of *ISO1*, therefore we had hoped that WGS of *HISR* line genomes might to point to a specific set of *P-*elements responsible for inducing strong GD. Unexpectedly, the *P*-element insertions were not confined to Chr2R, but rather were dispersed across the entire genomes of all *HISR* lines (Fig. 4A), seemingly defying the genomic PCR genotyping and WGS-SNP profiling that indicated sufficient backcrossing to favor mostly the *ISO1* genetic background (Fig. 4-S1).

**Figure 4.**
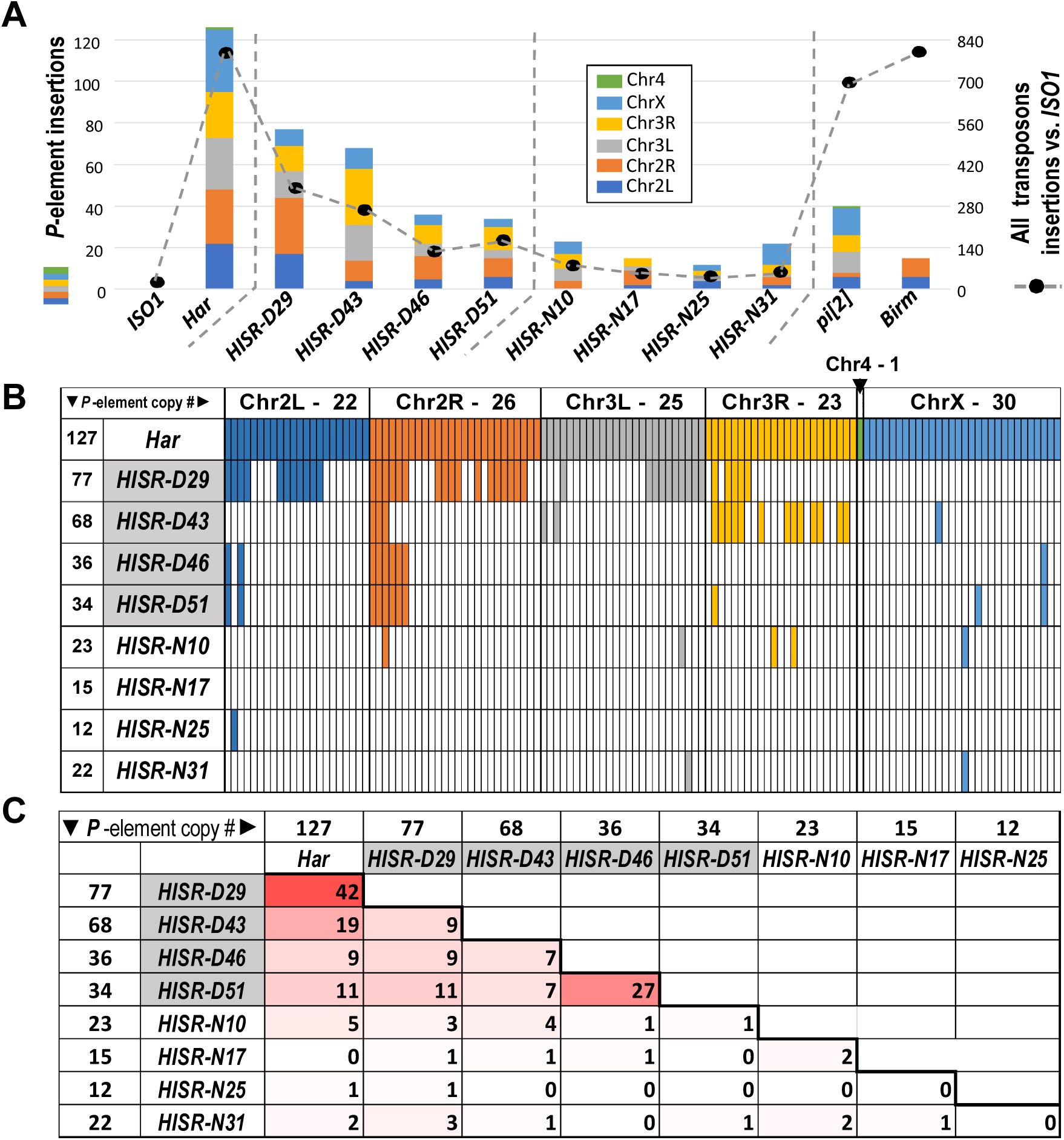
*P-elements* are mobilized *de novo* during generation of *HISR* lines. (A) TIDAL program counts of novel *P-*element insertions, left Y-axis and colored bars. Right Y-axis, black dots and dashed line are the total distinct transposon insertions in the unique-mapping portion of genome. (B) Lineage analysis of the *Har P*-elements retained in the *HISR* lines, colored by the major chromosomal segments. (C) Comparison of shared *P*-elements between *Har* and *HISR* lines, with total number of *P*-elements called by TIDAL in the top row and first column. Color shade reflects degree of shared *P*-elements between the two strains being compared.

To explore this conundrum, we examined how many of the original *P*-elements in the *Har* genome were conserved in the *HISR* lines’ genomes (Fig. 4B). As expected for *HISR-D29* whose *P*-element copy numbers was closest to *Har*, this line conserved the highest share of parental *Har P*-elements compared to other *HISR* lines. However, there were also 35 novel *P*-element insertions (∼45%) in *HISR-D29* absent from *Har*. Surprisingly, the vast majority of the *P*-element insertions across all *HISR* lines were also *de novo P-*element insertions (Fig. 4C), with each line clearing out nearly all parental *Har P*-element insertions and developing unique landscapes of *P-*element insertions. These data suggest that during the course of stabilizing the *HISR* lines, there were bursts of new *P*-element transpositions resulting in novel transposon landscapes that are completely distinct from the parental *Har* genome.

Although this dispersion of *de novo P*-elements in *HISR* lines’ genomes stymied our goal to pinpoint a particular *Har* locus strongly inducing GD, we next cloned and sequenced genomic PCR amplicons of all *P*-elements from the various *P*-element-containing strains. By using a single oligonucleotide that primes from both the 5’ and 3’ Terminal Inverted Repeats (TIRs), we amplified full-length *P*-elements as well as several additional truncation variants (Fig. 5A) that have been missed in other genomic PCR assays using internal primers (Wakisaka et al., 2017). The most abundant variant accounting for more *P*-element copies in *OreR-MOD* and *OreR-TK* strains compared to *Har* were the “*KP*” variant shown to encode a dominant negative protein that inhibits full-length *P*-transposase activity (Jackson et al., 1988; Simmons et al., 1990) (Fig. 5A–5C), thus explaining the innocuous accumulation of these *P*-element variants in these *OreR* strains. Full-length *P*-elements were also sequenced from *Har*, *pi*[*2*] and *Lerik-P* strains, but there were no sequence differences in these clones from the original full-length *P*-element sequence in GenBank that might suggest a superlative quality to the full-length *P*-element in these strong GD-inducing strains.

**Figure 5.**
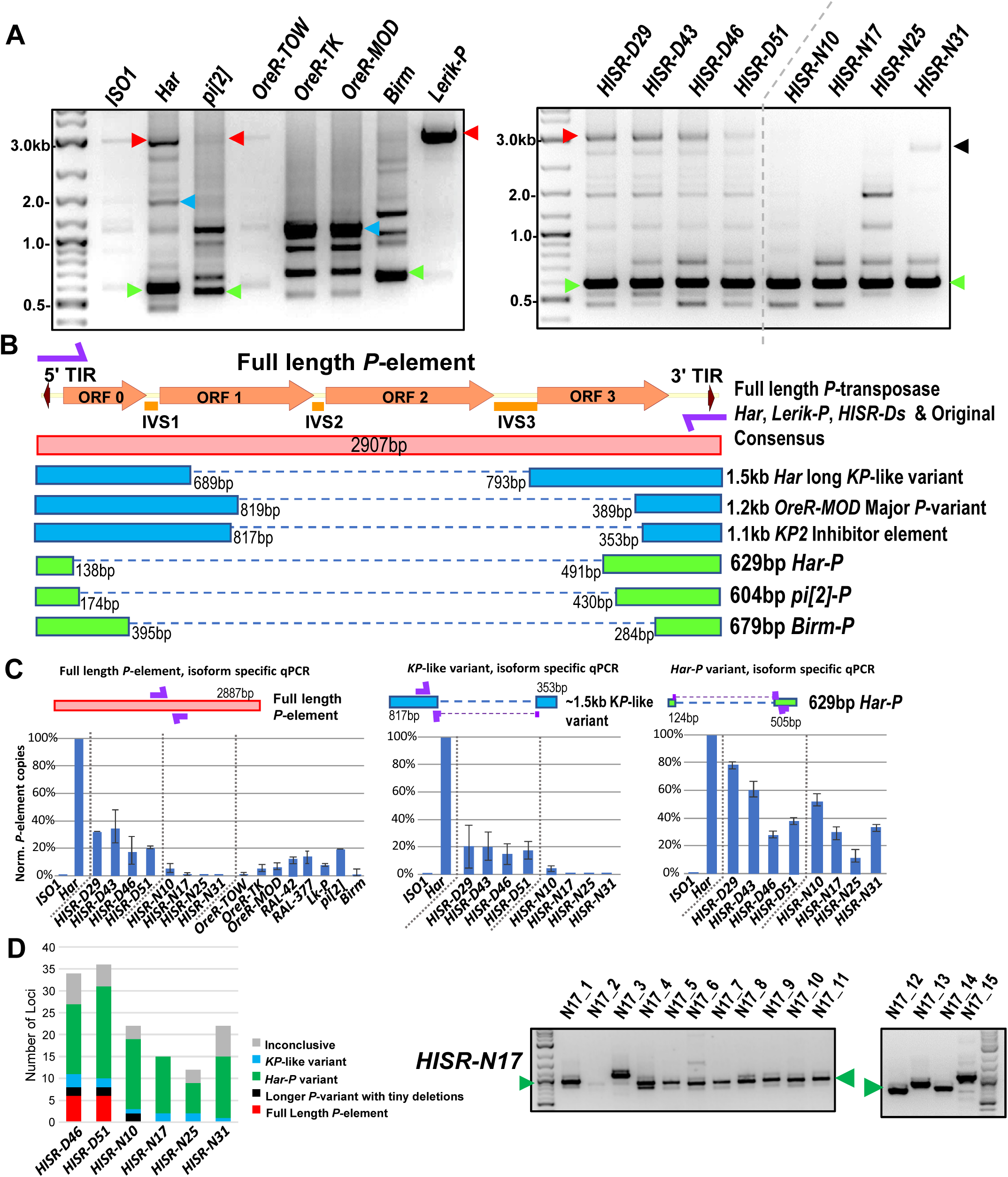
***Har-P* is a short and highly mobile *P-*element variant in strains used in *P*-element GD assays.** (A) *P*-element variant amplicons generated with TIR primers were cloned and sequenced as marked by colored arrows for the sequenced diagrams in (B). (B) Diagram of *P*-element variants cloned and sequenced from genomic PCR amplicons shown above. (C) Genomic qPCR quantifications of three *P*-element variants in *Harwich* and *Harwich*-derived *HISR* lines. Relative quantifications (in percentage) was calculated from ΔΔCt with *rp49* as reference gene. (D) Proportions of the *P-*element variants verified by locus-specific PCR from TIDAL predictions of all *HISR-N* and *HISR-D* strains with <40 *P-*element insertions. The gel for *HISR-N17* is on the right, while remaining gels are in Fig. 5-S1.

Interestingly, we sequenced a short ∼630bp *P*-element variant nearly-identical in *Har*, *pi*[*2*], and *Birm* strains, which only retains ∼130bp of the 5’ end and ∼500bp of the 3’ end of the *P*-element (Fig. 5A, 5B). By retaining functional TIRs, these short elements can still be detected by TIDAL in WGS, can mobilize during crosses with the *pi*[*2*] strain (Bingham et al., 1982; Mullins et al., 1989; O’Hare and Rubin, 1983), and assembles *in vitro* with the *P*-transposase tetramer complex >100X more efficiently than the full-length *P*-element (Tang et al., 2007). In all *HISR-D* lines that retain strong GD induction, we detected this short *P*-element variant and the full-length *P*-element encoding *P*-transposase, whereas the *HISR-N* lines retained the short variant but appeared to have lost the full-length *P*-element (Fig. 5A, right panel). With the smaller number of TIDAL-predicted *P-*element insertions in *HISR-N* lines, we confirmed by locus-specific PCR the absence of full-length *P*-elements and that the majority of *P*-element insertions (∼55-95%) were these *de novo* short *P*-element insertions (Fig. 5D and 5-S1). We name this short variant “*Har-Ps*” (*Harwich P’s*) in homage to Harpies, highly mobile hybrid bird-human creatures from the Greek mythological stories of the Argonauts.

### Restoring GD when Har-P is crossed with P-transposase expressed only in the germline

We hypothesized that *Har-Ps* combined with *P*-transposase from full-length *P*-elements could be the drivers of strong GD induction from *pi*[*2*], *Har*, and *HISR-D* strains. To test this hypothesis, we used negative-control *yw*-background females that lack *P*-transposase and transgenic *H{CP}3* females that only express *P*-transposase in the germline (Simmons et al., 2002b) in crosses with males that either lack *Har-P* copies (*ISO1, Lerik-P, OreR-MOD*) or contain many *Har-P* copies (*Har, HISR-N’s, Birm*)(Fig. 6A). GD induction was only restored in the F1 daughters of this cross in strains with many *Har-Ps* (Fig. 6B). To avoid silencing of *P*-transposase by maternal *P*-element piRNAs in these strains, these crosses specifically used males that should only contribute paternal chromatin without contributing piRNAs(Fig. 6A). Notably, the *KP*-length and full-length *P*-elements in *OreR-MOD* and *Lerik-P*, respectively, did not restore GD (Fig. 6B, right most bars of left graph). These results suggest *P*-transposase act upon *Har-P* loci rather than longer *P*-element variants to induce GD and support the observation for *Har-P* loci making up the majority of the *de novo P-*element insertions in *HISR-N* lines (Fig. 6C). Our data is also consistent with a previous study showing that *P*-transposase assembles much more efficiently *in vitro* on short *P*-elements compared to the full-length *P*-element (Tang et al., 2007).

**Figure 6.**
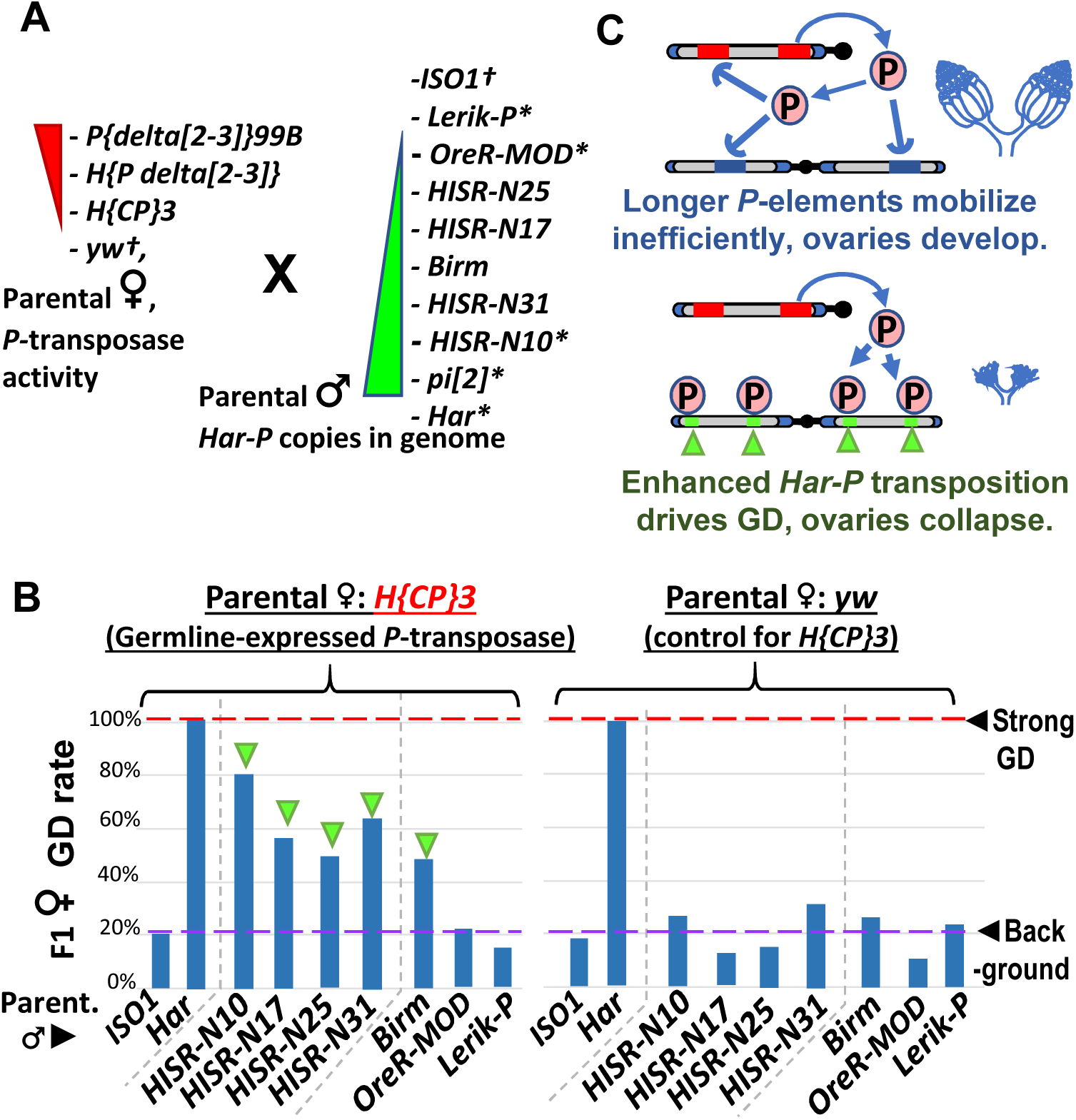
In absence of piRNA silencing, *Har-P* crossed with *P*-transposase restores severe GD. (A) Strains for testing *Har-P* genetic interaction with *P*-transposase activity. *-strains with piRNA silencing; † strains lacking any *P*-elements. (B) The *H{CP}3* strain’s moderately-expressed maternal dose of *P*-transposase crossed with paternal *HISR-N* strains and *Birm* strain restores GD in F1 daughters, but strains with longer and full-length *P*-elements like *OreR-MOD* and *Lerik-P* lack the GD phenotype. (C) Model for *P*-transposase mobilizing *Har-Ps* to cause catastrophic transposition.

We noticed that GD severtiy in crossing *HISR-N* with *H{CP}-3* strains was not completely penetrant like GD assays with the parental *Har* (Fig.6B versus Fig 1B) because *Har* contributes both multiple copies of full-length *P*-elements and *Har-P* loci versus the single copy of the natural *P*-element transgene in *H{CP}-3* (Simmons et al., 2002b). In addition, natural *P*-element translation is inhibited by strong somatic splicing inhibition of the native *P*-element’s third intron (IVS3) containing a premature stop codon and only inefficient splicing in the *Drosophila* germline that is further suppressed by piRNAs (Siebel et al., 1994; Teixeira et al., 2017). We also confirmed that IVS3 intron splicing was the main alteration that increased *P*-element expression in ovaries from a dysgenic cross between *Har* and *ISO1*, whereas Open Reading Frame (ORF) parts of the P-element transcript are only modestly increased (Fig. 6-S1A). We believe this sufficient expression of *P*-transposase promotes the preferred mobilization *Har-P* short variants in dysgenic cross ovaries, but the cut-and-paste transposition mechanism of *P*-transposase should theoretically conserve the total copy number of *P*-elements. By using digital droplet PCR to precisely quantity total *P*-element copy numbers, we confirmed that total *P*-element copy numbers were stable across ovaries of daughters from two sets of dysgenic and non-dysgenic crosses (Fig. 6-S1B).

### Somatic expression of P-transpose with Har-P’s causes pupal lethality

To test whether a stronger expressing P-transposase transgene could induce the complete GD in crosses with *HISR-N* lines, we turned to the *delta*[*2-3*] *P*-transposase transgenes that lack the IVS3 intron to enable strong somatic and germline *P*-transposase activity (Robertson et al., 1988). When we crossed two different *delta*[*2-3*] female strains to males of *HISR-N17*, *-N25*, and *-N31* which lack *P*-element piRNA expression but have *Har-Ps,* we were unable to assay GD because of extensive pupal lethality (Fig. 7A). We also confirmed extensive pupal lethality in crosses between *delta*[*2-3*] and the *Birm* strain (Fig. 7A) as previously described (Engels et al., 1987; Simmons et al., 1987). Since we also detected very short *P* variants in *Birm* that are similar to *Har-P* (Fig. 5A, 5B) we conclude that somatically expressed *P*-transposase acting only on the *Har-Ps* in *Birm*, *HISR-N17*, *-N25*, and *-N31* is sufficient to disrupt pupal development.

**Figure 7.**
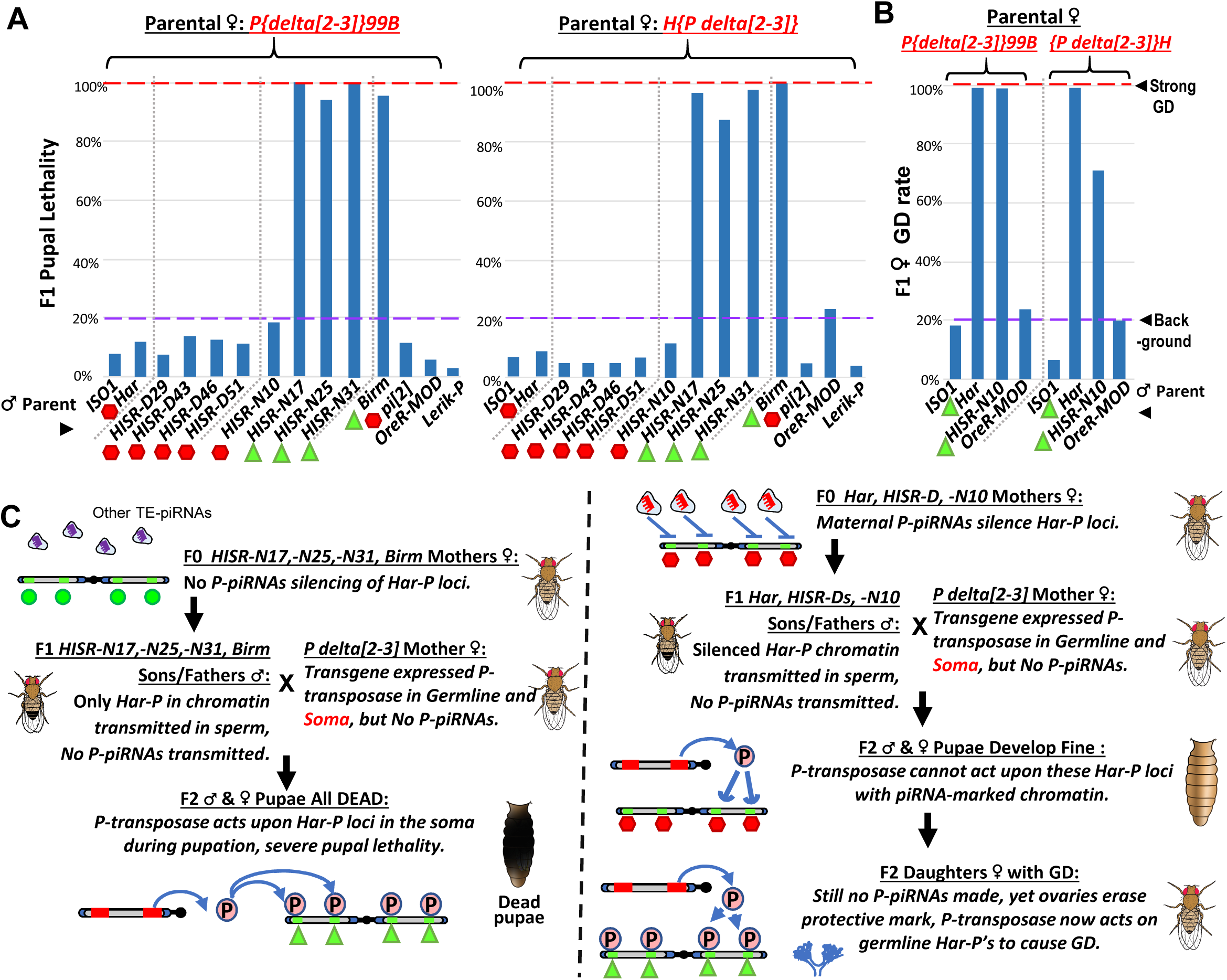
Somatic expression of *P*-transposase triggers pupal lethality with *Har-P* loci that are not silenced by *P-element* piRNAs. (A) Green triangles denote crosses showing pupal lethality from stronger somatic expression of *P*-transposase acting on *Har-Ps* in *HISR-N* and *Birm* strains lacking *P*-element piRNAs. Red hexagons denote crosses with strains expressing *P*-element piRNAs that suppress pupal lethality through a paternally transmitted epigenetic imprint. (B) The paternal *P*-element piRNA imprinting on *Har-Ps* in *Har* and *HISR-N10* cannot suppress GD in F1 daughters, as marked by green triangles. The longer *P* variants in *OreR-MOD* do not result in GD with the *delta2-3 P-*transposase. (C) Revised *P* Dysgenesis paradigm proposing a paternally-transmitted piRNA-directed epigenetic mark that resists *P*-transposase activity in the soma, but this mark is erased during oogenesis.

Unexpectedly, the pupal lethality was not observed when *delta*[*2-3*] females were crossed with *Har-P*-containing males that also expressed *P*-element piRNAs, such as *Har*, *pi*[*2*], four *HISR-D* lines, and *HISR-N10* (Fig. 7A). These hybrid F1 progeny developed into adults, but the adult females of *Har* and *HISR-N10* hybrids with *delta*[*2-3*] still exhibited absolute GD (Fig. 7B). These data suggest that *P*-element piRNAs impart a paternally-transmitted imprint on *Har-P* loci that resists mobilization with somatically-expressed *P*-transposase and enables development to adulthood. However, this imprint is either erased in ovaries or insufficient to prevent ovarian GD. Finally, the notable *P*-element piRNA pattern of *HISR-N10* perfectly matches the *Har-P* structure since many internal piRNAs are absent (Fig. 3B), but overall *P*-element piRNAs in *HISR-N10* are equivalent to *Har* and *HISR-D* lines (Fig. 3C), and therefore are sufficient to repress *Har-P’s* epigenetically from being mobilized in the soma by the *delta2-3 P*-transposase.

## DISCUSSION

After a *Drosophila* strain has silenced an invading transposon through the Piwi/piRNA pathway, the neutered transposon will naturally decay into various truncations that are presumed to be neutral or even beneficial to host fitness (Kelleher, 2016), such as natural *KP2* truncation variants that inhibit *P*-element transposition (Jackson et al., 1988; Simmons et al., 1990). However, we discovered one such truncation we call *Har-P* via our unbiased genetic and molecular approach that can actually be detrimental to the host. Since *P*-transposase assembles *in vitro* much more efficiently on very short natural *P*-element variants (Tang et al., 2007), we propose a new model for catastrophic *P*-element transposition in strong GD inducer strains like *pi*[*2*] and *Har (*Fig 6C). When a *P*-element truncates to a ∼630bp *Har-P* variant, this non-autonomous variant dominates as the main mobilizing *P*-element during a dysgenic cross to induce strong GD. Therefore previous studies examining GD variability across other *Drosophila* strains and isolates may now be explained by whether these genomes contain both full length and very short *P*-elements (Bergman et al., 2017; Kozeretska et al., 2018; Ronsseray et al., 1989; Srivastav and Kelleher, 2017; Wakisaka et al., 2017; Yoshitake et al., 2018).

Although our future goal will be to determine which specific epigenetic marks are deposited at full length *P-elements* and *Har-P’s* by piRNAs, we believe a chromatin mark resisting *P*-transposase activity is more likely than somatic piRNAs or siRNAs (Chung et al., 2008; Ghildiyal et al., 2008; Kawamura et al., 2008) silencing the *delta2-3 P*-transposase in our pupal lethality crosses because we confirmed robust *P-* transposase mRNA expression regardless of the expression of *P-element* piRNAs (Fig. 7-S1A). A second future goal will be to generate transgenic flies with single or multiple synthetic *Har-P* copies to determine the precise dosage of *Har-P’s* that would trigger GD or pupal lethality. However, in addition to copy number, genomic location may also influence host tolerance of *Har-P’s*, because we observed a significant rescue of viable pupae in crosses between *delta2-3 P*-transposase and a derivative strain of *HISR-N17* with *Har-P’s* only on Chromosome 3 with six *P-elements*, while no pupae survived with *delta2-3 P-*transposase and *Har-P’s* on Chromosome 2 with nine *P-elements* (Fig. 7-S1B).

The *Drosophila P*-element system of hybrid GD mainly affects female sterility and requires maternally contributed *P*-element piRNAs to propagate transgenerational *P*-element silencing in daughters via trimethylation of histone H3 lysine 9 (H3K9me3) (Josse et al., 2007; Le Thomas et al., 2014). Although previous studies of dysgenic crosses focused on complete GD in females (Bingham et al., 1982; Brennecke et al., 2008; Khurana et al., 2011; Rubin et al., 1982), sons respond differently because they are fertile despite presumed somatic *P*-element excision (Wei et al., 1991). Mouse piRNAs bound by MIWI2 direct the re-establishment of DNA methylation marks on transposons like *L1* and *IAP* (Aravin et al., 2008), which may propagate in sperm, but *Drosophila* lacking DNA methylation and previous studies of P-M hybrid dysgenesis never considered a paternal imprint on *P*-elements that our findings now suggest is being propagated (Fig. 7C). Similar to other metazoans, *Drosophila* sperm also undergoes histone exchange with protamines (ProtB), with little contribution of paternal cytoplasm (Rathke et al., 2007). However, recent data do support the retention of some H3K9me3 in sperm (Yamaguchi et al., 2018), which might underlie the paternal imprint of piRNA-silencing of *P*-elements that will be investigated further in future studies.

This interplay between the truncated *Har-Ps* and full-length *P*-element DNA transposon resembles other examples in nature, such as the extreme proliferation of MITEs (miniature inverted repeat transposable element) in rice scavenging other transposases to mobilize (Yang et al., 2009), and short mammalian SINEs retrotransposons taking advantage of the transposition machinery of longer LINEs, since SINEs persist in much greater numbers than their longer LINE counterparts (Hancks and Kazazian, 2012). However, while the full impact of MITEs and SINEs on organism development is still obscure, our study indicates that *Har-Ps* combined with the *P*-transposase to trigger transposition events so efficiently to be detrimental to ovarian and pupal development. Notwithstanding, the high efficiency of *Har-P* mobilization *by P*-transposase may also be engineered into a new generation of transposon-based mutagenesis approaches.

## MATERIALS AND METHODS

### Fly strains

All strains were maintained on standard cornmeal medium at 22°C. Because the *ISO1*(BDSC#2057) stock had accumulated >180 new transposon insertions relative to the original stock sequenced in the Berkeley *Drosophila* genome project (Adams et al., 2000; Rahman et al., 2015), we obtained the *ISO1* strain from Susan Celniker’s lab (*ISO1-SC*). The *Har* strain was obtained from (*Har-WET*) was obtained from the William Theurkauf’s lab (Khurana et al., 2011). Three *Oregon-R* strains were obtained from Terry Orr-Weaver’s lab, *OreR-TOW*, *OreR*-TK (Kaufman, BDSC#2376) and *OreR-MOD* (BDSC#25211). The *Lerik-P* strain was obtained from Stephane Ronserray’s lab (Josse et al., 2007; Marin et al., 2000). All the following strains were also directly obtained from the BDSC – *RAL-42* (#28127), *RAL-377* (#28186), *pi*[*2*] (#2384), *y*[*1*] *w*[*67c23*]*; H{w*[*+mC*]*=hsp/CP}3* (#64160), *Birmingham; Sb*[*1*]*/TM6* (#2539), *w*[*\**]*; wg*[*Sp-1*]*/CyO; ry*[*506*] *Sb*[*1*] *P{ry*[*+t7.2*]*=Delta2-3}99B/TM6B, Tb*[*+*] (#3629), and *y*[*1*] *w*[*67c23*]*; H{w*[*+mC*]*=w*[*+*]*. Delta2-3.M}6* (#64161). *Sp/CyO;TM6b/Sb* was obtained from Michael Rosbash’s lab.

### Crosses, gonadal dysgenesis and pupal lethality assays

All crosses were set up with 3-5 virgin females and 2-4 young males per replicate on standard cornmeal medium at 25°C and parents were purged after 5 days of egg laying (Srivastav and Kelleher, 2017). For GD assays, F1 females aged to 4-5 days at 25°C were examined for GD using food dye and GD % shown is average of 3 replicate crosses with total minimum of 100 F1 females assayed (Simmons et al., 2007). Somatic pupal lethality was recorded by counting dead (uneclosed) and empty pupae (eclosed) 6 days after first eclosion was observed in respective control cross (*P{delta*[*2-3*]*}99B x ISO1* or *H{P delta*[*2-3*]*} x ISO1*) (Engels et al., 1987). Pupal lethality percentage shown is average of two or more replicate crosses that obtained at least total of 50 F1 pupae each.

### Crossing scheme to generate *HISR* lines

The detailed crossing scheme is illustrated in Fig. 2-S1. After a first cross between virgin *Har* females and *ISO1* males, three more backcrosses of virgin *Har/ISO1* hybrid progeny females mated to *ISO1* males were performed and following the progeny with red eyes to select for the *Har* segment segregating with the *cn*, *bw, sp*, alleles on Chromosome 2R. We hoped that a particular set of *P*-elements that drive strong GD induction would co-segregate with red eye color. We then performed a ‘Validation Cross’ with the F4 hybrid males individually mated to *ISO1* females. We screened >100 individual groups of F4 males for their GD induction, where the early-hatching 3-day old daughters were screened via the squash assay for 100% GD. Only the F5 vials showing 100% GD from F4 males crossed to *ISO1* females were kept, and then were allowed to age and self-crossed and propagated in 11 more generations to attempt to create recombining-inbred-lines (RILs).

Selecting only flies with red eyes required purging any flies emerging with the “white” eyes of *ISO1* and discarding many vials that failed to generate progeny due to genotoxic collapse from inability to silence *P*-element transposition. At the F16 stage, <u>H</u>ar/<u>I</u>SO1 <u>S</u>elfed <u>R</u>ed (*HISR)* lines males were rescreened in a Validation Cross with *ISO1* females, this time keeping lines that still caused 100% GD and designated as *HISR-D* (Dysgenic) lines. We also selected additional lines that had now lost GD and allowing for >50% of females to generate egg chambers, and these were designated *HISR-N* (Non-dysgenic) lines. We performed 2 rounds of single-sibling pair mating to further inbreed these lines in an attempt to stabilize the genotypes, and we maintained 4 lines of each *HISR-D* and *HISR-N* for true propagation of just the red or cinnabar eyes and speck phenotype.

### Genomic DNA extraction, PCR, quantitative-PCR and Droplet Digital PCR

Genomic DNA was prepared from 10 young female flies by homogenizing tissues with plastic pestle in 300µL Lysis buffer (10mM Tris pH-8.0, 100mM NaCl, 10mM EDTA, 0.5% SDS, and Proteinase K at 50µg/ml) and incubated at 65°C overnight followed by treatment of RNase A at 100 µg/ml at 37°C for 30 mins. 200µL of 0.5M NaCl was added followed by one volume of Phenol:CHCl_3_:IAA (at 25:24:1) and spun at 14,000 rpm for 10 minutes to isolate DNA in aqueous phase. Aqueous phase was extracted again with one volume of CHCl_3_:IAA (at 24:1) and supplemented with one volume of 5M LiCl and incubated at −20°C and then spun at 15,000 rpm for 15 mins to precipitate RNA. Supernatant was isolated and supplemented with 2 volumes of 100% ethanol and incubated in −20°C for 2 hours and then spun at 15,000 rpm for 20 mins. DNA pellet was washed with chilled 70% ethanol and dissolved in nuclease free water. DNA integrity checked (>10kb) by running 1 µg on 1% agarose gel with EtBr.

Genomic PCR reactions to characterize *P*-element structural variation were set up in 30µL reactions of 1X NEB GC buffer, 300µM dNTPs, 0.5M Betaine, 2.5mM MgCl_2_, 0.25µM of IR primer (Rasmusson et al., 1993), 1µL of Phusion polymerase and 50ng of genomic DNA and cycled at 94°C for 1 min, 62°C for 2 mins, 72°C for 4 mins for 27 cycles and followed by 72°C for 15 minutes. Genomic PCR reactions to characterize *P*-element structural variation in *HISR* lines, predicted by TIDAL were also set up similarly using *P*-element insertion locus specific primers. Genomic PCR reactions for genotyping of *HISR-N* lines were set up similarly but cycled at 94°C for 30 sec, 60°C for 15 sec, 72°C for 30 sec for 27 cycles and followed by 72°C for 5 minutes.

Genomic qPCR experiments were performed in three biological replicates with two 20µL technical reactions replicates each, using Luna Mastermix (NEB), primers at 0.5µM and 20ng of genomic DNA per reaction in real time quantitative PCR. *P*-element load was calculated from 2^(-ΔΔCt) normalized to *Har* at 100% and ΔCt from *RP49*. All primers used for are listed in Table S1.

For the Droplet Digital PCR (ddPCR), we utilized the Evagreen Mastermix (Biorad) and conducted on a QX500 ddPCR machine with manual setting of droplet signal thresholds. 10-15 pairs of ovaries and corresponding carcass from 4-5 day old F1 females was dissected from dysgenic and non-dysgenic crosses of *Har* and *HISR-D51* with *ISO1* strain at 18°C. DNA was extracted from the ovaries and carcass and quantified using Qubit 2.0 Fluorometer. Digital PCR probe assays were conducted in 40µL droplet reactions, generated from 25µL digital PCR reaction and 70µL droplet oil each. 25µL digital PCR reactions were set up with BioRad ddPCR probe supermix, *P*-element7a (FAM) and rp49 (HEX) probes each at 250nM and 200pg of DNA. Reactions were cycled at 95°C for 10 mins followed by 95°C for 30s and 58°C for 1 min for 40 cycles, and 98°C for 10 mins. Copies/µL values were extracted from QuantaSoft (BioRad) software and *P*-element copies per genome were calculated normalized to rp49.

### *P*-element amplicon cloning and sequencing

*P*-elements amplified from IR PCR were purified from 1% agarose gel using QIAquick Gel extraction kit and cloned into pCR4-TOPO vector using Zero Blunt TOPO PCR Cloning Kit at RT, followed by transformation of chemically competent DH10β cells, which were then grown on LB plates with 0.05mg/ml Kanamycin overnight. 5-10 colonies were screened by PCR and two colonies positive for *P*-element cloned were chosen for plasmid mini-prep and sequenced using M13 forward and reverse primers for all variants in addition to internal primers to complete the sequencing of full-length *P*-elements.

### Whole genome sequencing, SNP profile analysis, and TIDAL analysis

Genomic DNA libraries were prepared using NEB Ultra II FS kit E7805. Briefly, 500ng of genomic DNA (>10kb) was fragmented at 37°C for 12 minutes, followed by adaptor ligation and loop excision according to kit manual protocol. Size selection was performed with two rounds of AmpureXP beads addition to select for insert size 150-250bp as per kit manual. Library PCR amplification was also carried out as per manual instructions for 6 cycles and purified using one round of AmpureXP beads addition at 0.9X volume. Individual barcoded libraries were quantified on NanoDrop and each diluted to 2nM and then pooled to produce equimolar concentration.

Whole genome sequencing was performed on an Illumina NextSeq 500 with paired-end reads of 75bp x 75bp in the Rosbash lab at Brandeis University. Reads were demultiplexed and trimmed by Trimmomatic to remove low quality bases, and then reads were analyzed by the TIDAL program (Rahman et al., 2015). TIDAL outputs were sorted for *P*-element insertions and the insertion coordinates were compared across the *HISR* lines using SQL queries in MS-Access. To calculate the Single Nucleotide Polymorphism (SNP) profiles, paired-end reads were mapped to the Dm6 *ISO1* genome with "BWA MEM"(Li and Durbin, 2010) using default parameters. PCR duplicates are removed with Picard and SNPs are called with GATK HaplotypeCaller (Danecek et al., 2011; DePristo et al., 2011; McKenna et al., 2010). We then generated the nucleotide distribution for each SNP to ensure that there are at least 20 reads supporting each SNP. Then we created a unified SNP list by using the union of SNPs from all libraries and carefully noted if each SNP is present in each library. The SNP counts were binned by 5kb segments and converted into a graphical representation as differences between the reference genome and strain/line in Fig. 4-S1.

### Ovary small RNA sequencing and analysis

To remove the 2S rRNA from *Drosophila* ovaries, we adapted a protocol from our previous Q-sepharose beads matrix technique (Lau et al., 2009). About 50 ovaries per parental *Har* and *ISO1* strains and *HISR* lines were dissected from young adult females. Ovaries were then lysed in ice cold 500ul Elution Buffer (20mM Hepes pH 7.9 (with KOH), 10% glycerol, 400 mM KOAc, 0.2 mM EDTA, 1.5 mM MgCl2, 1.0 mM DTT, 1X Roche Complete EDTA-free Protease Inhibitor Cocktail) using 1 freeze-thaw cycle and pulverizing with a blue plastic pestle. A 1.5ml aliquot of Q-Sepharose FF matrix suspension was washed 1X in water, then 3X in Elution buffer, then incubated for 10 minutes with the ovaries lysate with occasional agitation in cold room. Ribosomal RNA gets bound by the Q-sepharose, while small RNA RNPs remains in the elution buffer. Elution buffer was removed and then subjected to small RNA extraction with the Tri-reagent protocol. The precipitated small RNAs where then converted into Illumina libraries using the NEBNext Small RNA Library Construction kit. One modification we employed during the overnight linker ligation is to supplement the reactions to 12.5% PEG 8000 to reduce the potential sequence biases from T4 RNA ligase activity.

Small RNA libraries were sequenced as 75bp single end reads on the NextSeq550. Adapters for the small RNA libraries were removed with CutAdapt and then mapped to the *Drosophila* transposon consensus sequences from RepBase and Flybase using Bowtie v1 with 1 mismatch and R plotting scripts as applied in our previous published studies on *Drosophila* piRNAs (Clark et al., 2017; Sytnikova et al., 2014).

### RT-qPCR analysis of *P*-element expression in gonadal dysgenesis and pupal lethality

For this assay, 5-10 pairs of ovaries were dissected from 3-5 day old F1 females of dysgenic and non-dysgenic cross with *Har* and *ISO1*, as well as with *Har* and *HISR-D46.* RNA was extracted from such ovaries and integrity checked by running 1µg RNA at 2% Agarose II gel (Fischer BioReagents). 3µg was reverse transcribed using Protoscript RT enzyme (NEB) as per manufacturer’s protocol and negative RT control was carried out similarly without RT enzyme. 50ng of cDNA was used for setting up rp49 PCR reactions (as described above) from RT and corresponding negative RT reactions to evaluate DNA contamination. qPCR reactions for *P*-element ORF2, ORF3, IVS3 were also carried out as genomic qPCR reactions with 20ng cDNA input and ΔCt were calculated similarly using rp49 RNA levels.

In the RT-qPCR analysis of *H{P delta*[*2-3*]*}* gene expression in gonadal dysgenesis and pupal lethality, 5-10 pairs of ovaries and corresponding carcass were dissected from F1 females of pupal lethality crosses conducted at 18°C. RNA extraction, reverse transcription, PCR and qPCR reactions were carried out similarly as above. ΔCt were calculated similarly using rp49 RNA levels. Fold change values were obtained from normalizing F1 carcass *P*-element RNA levels to *H{P delta*[*2-3*]*}* carcass and F1 ovary P-element RNA levels were normalized to *H{P delta*[*2-3*]*}* carcass in Fig 7-S1.

### Isolation of HISR-N17 autosomes for modulating *Har-P* genomic dosage

HISR-N17 autosomes were isolated first by crossing virgin HISR-N17 females with Sp/CyO;TM6b/Sb stock males and using virgin F1 females with CyO and TM6 to cross again with Sp/CyO;TM6b/Sb. F2 males with either HISRN-17 Chr2 or HISR-N Chr3 were crossed to virgin *H{P delta*[*2-3*]*}.* All crosses were performed in triplicates at 25°C. F3 pupal lethality was recorded on 16^th^ day of the *H{P delta*[*2-3*]*}* crosses.

## ACKNOWLEDGEMENTS

We thank W. Theurkauf, S. Celniker, S. Ronsseray, T. Orr-Weaver, and the BDSC (NIH grant P40OD018537) for fly strains; M. Rosbash and Brandeis University for deep sequencing and fly food; and D. Schwarz, R. McCrae, A. Grishok and D. Cifuentes for comments. SPS and NCL conceived and conducted the experiments, RR and QM provided bioinformatics analyses, and NCL wrote the paper. This work was supported by NIH grants R01-AG052465 and R21-HD088792 to NCL. Sequencing data is deposited in the NCBI SRA as Study #SRP178563.

## SUPPORTING ONLINE MATERIAL

**Figure 2-S1.**
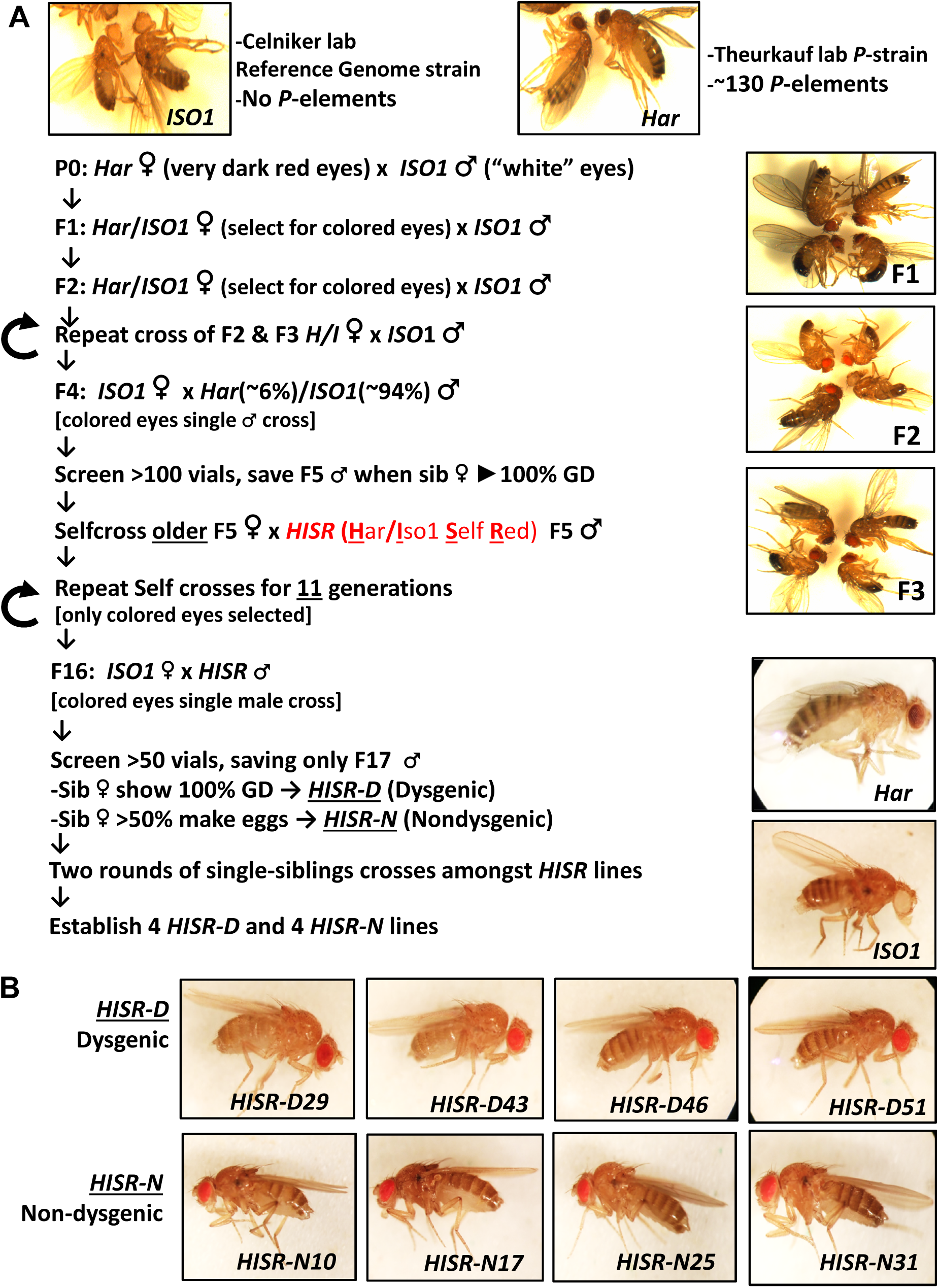
Detailed scheme for generating hybrid strains of reduced *Har* genome within the *ISO1* background. (A) Genetic crossing scheme to generate <u>H</u>ar/<u>I</u>so1 <u>S</u>elfed <u>R</u>ed (*HISR*) lines based on the tracking of eye color and retention of strong gonadal induction during a paternal cross to the *ISO1* strain. (B) Eight independently-selected lines that either retained or lost strong gonadal dysgenesis, with comparison to the parental lines above.

**Figure 2-S2.**
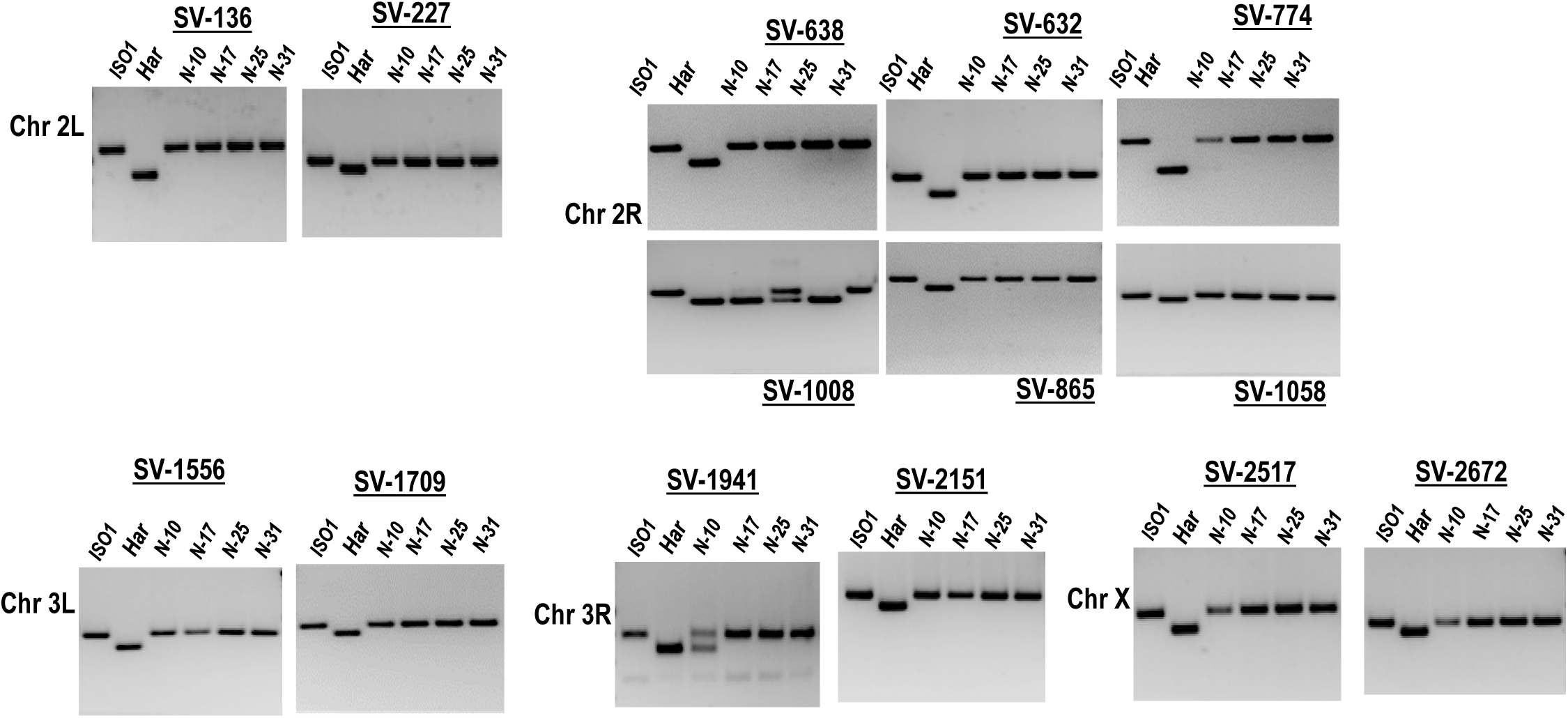
Genomic PCR genotyping of *HISR* strains. PCR amplicons of several loci in *HISR-N* strains are consistently homozygous for the *ISO1* allele. PCR conditions and primers are detailed in Table S1.

**Figure 3-S1.**
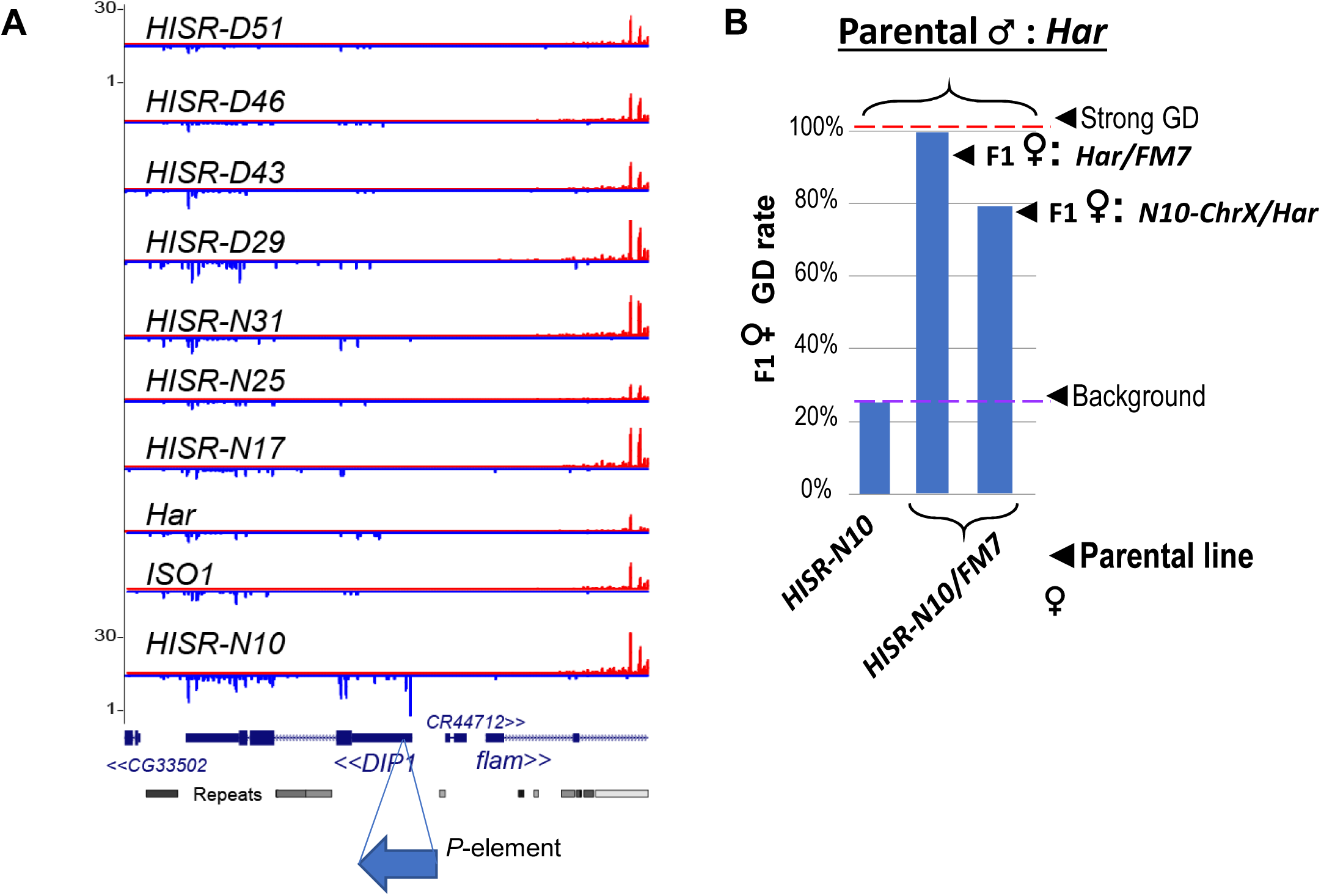
**Candidate locus for *P*-element directed piRNA production in *HISR-N10* ovaries.** (A) Genome browser plots of small RNAs mapping to the *DIP1* locus, adjacent to *Flamenco* piRNA locus, where only *HISR-N10* has a novel *P*-element insertion at this region that is causing increased piRNAs at this locus. (B) A parental female of *HISR-N10* with the *FM7* balancer chromosome for X are then crossed with *Har* males to test for GD repression as in Fig3C.

**Figure 4-S1.**
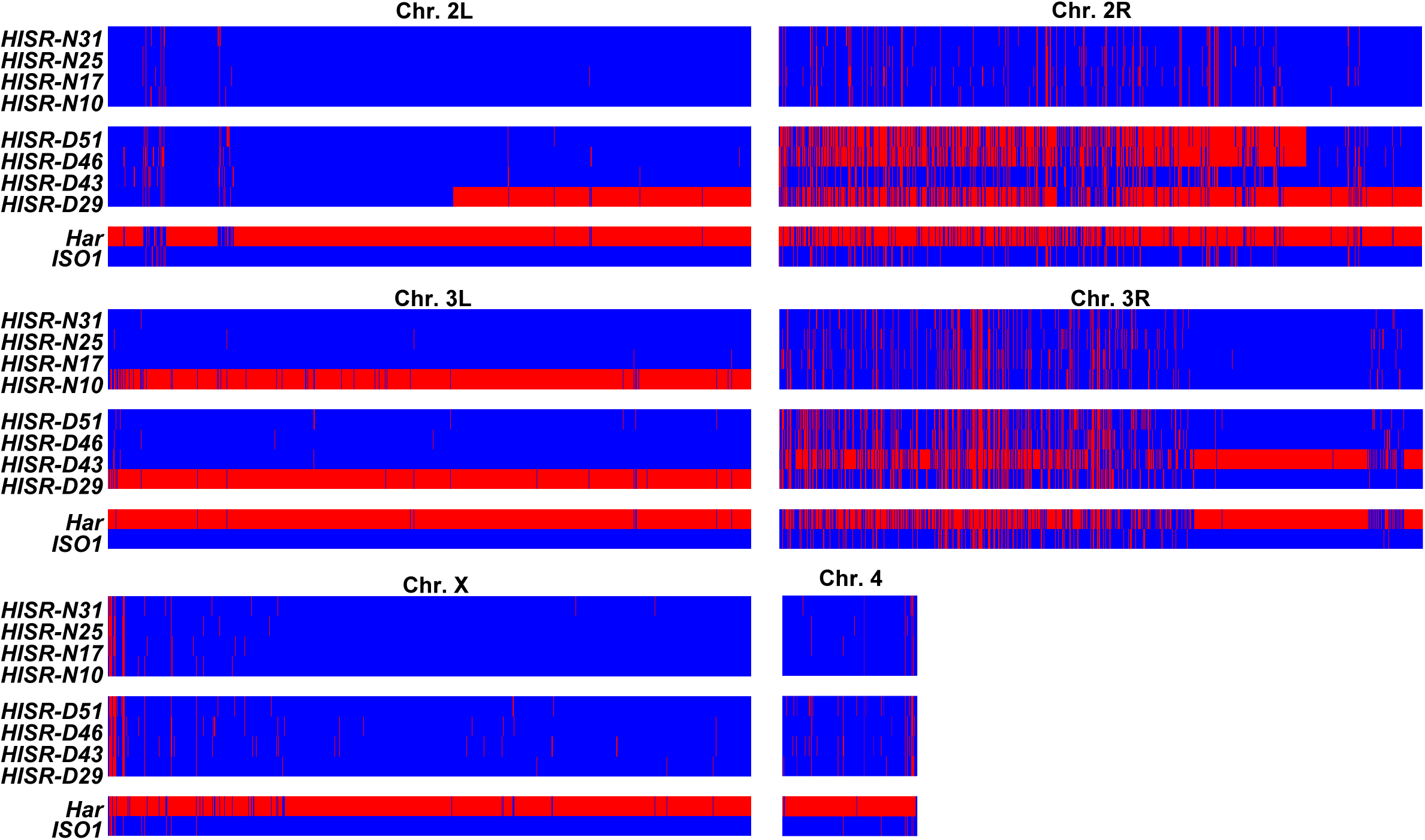
Genome-wide SNP profiling of *HISR* strains. Single Nucleotide Polymorphism (SNP) profiles comparisons between *HISR* strains and parental strains. For each 5kb interval, red marks a SNP distinct from the *ISO1* reference genome, whereas blue marks identity with reference genome sequence.

**Figure 5-S1.**
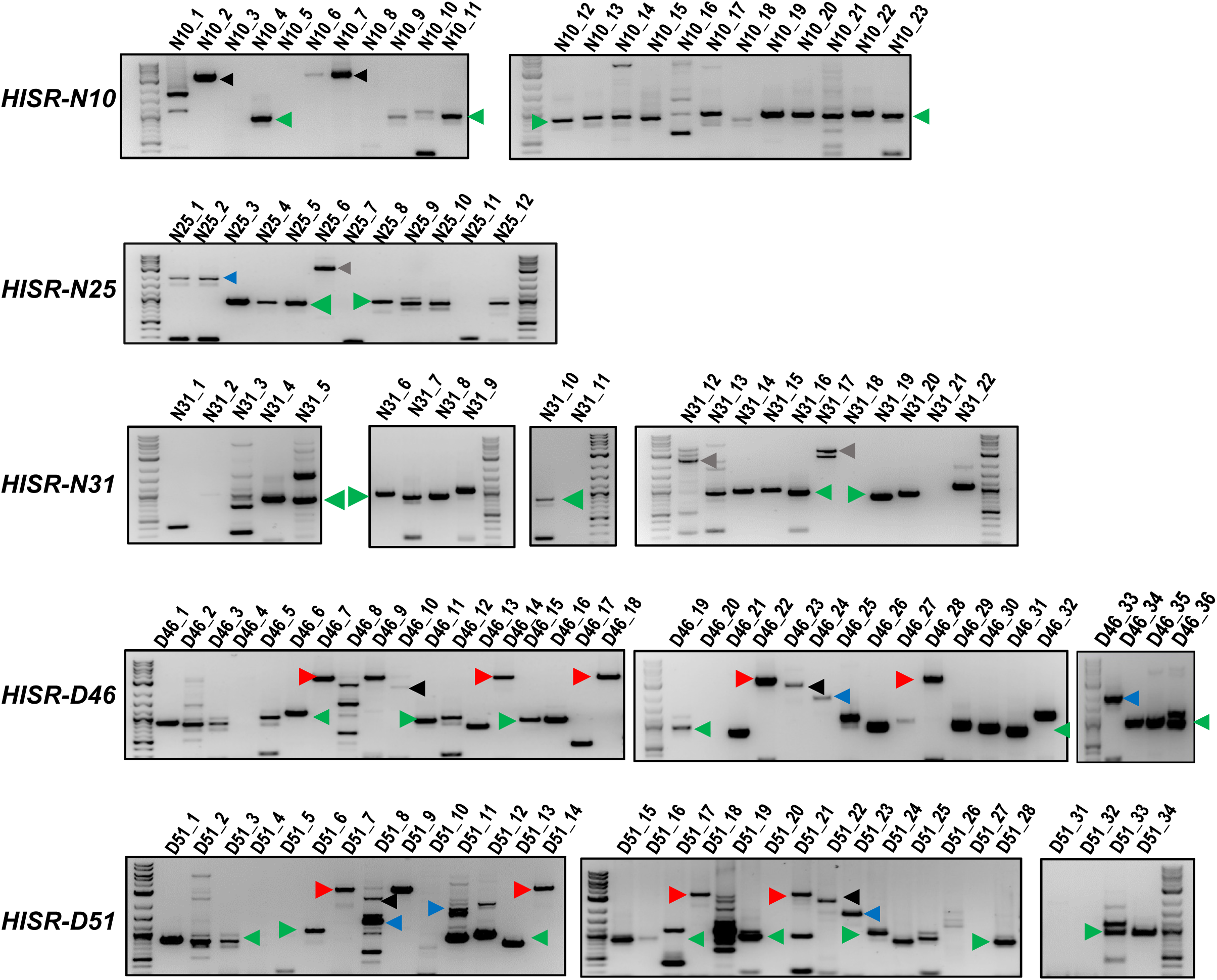
Genomic PCR amplifications of *P*-element insertion loci in *HISR-N* and *HISR-D* lines. Characterizing variant distributions among *P-element* insertions predicted in all *HISR-N and* two *HISR-D* genomes (*-D29* and *-D43* strains were omitted because >40 *P*-element insertions were predicted by TIDAL, Fig. 4). Loci number represent the predicted *P*-element insertion sites from TIDAL analysis of Illumina whole genome sequencing. Green arrowheads mark ∼0.6kb *Har-P* variants, blue arrowheads mark the likely *KP*-like variant, red arrowheads mark full-length *P*-elements, and black arrowheads mark full length or uncharacterized *P*-variants. Quantitation of variants proportions shown in Fig. 5D. The similar patterns of amplicons between *–D46* and *-D51* strains is expected since a majority of the new *P*-element insertions are shared between these two strains (Fig. 4C).

**Figure 6-S1.**
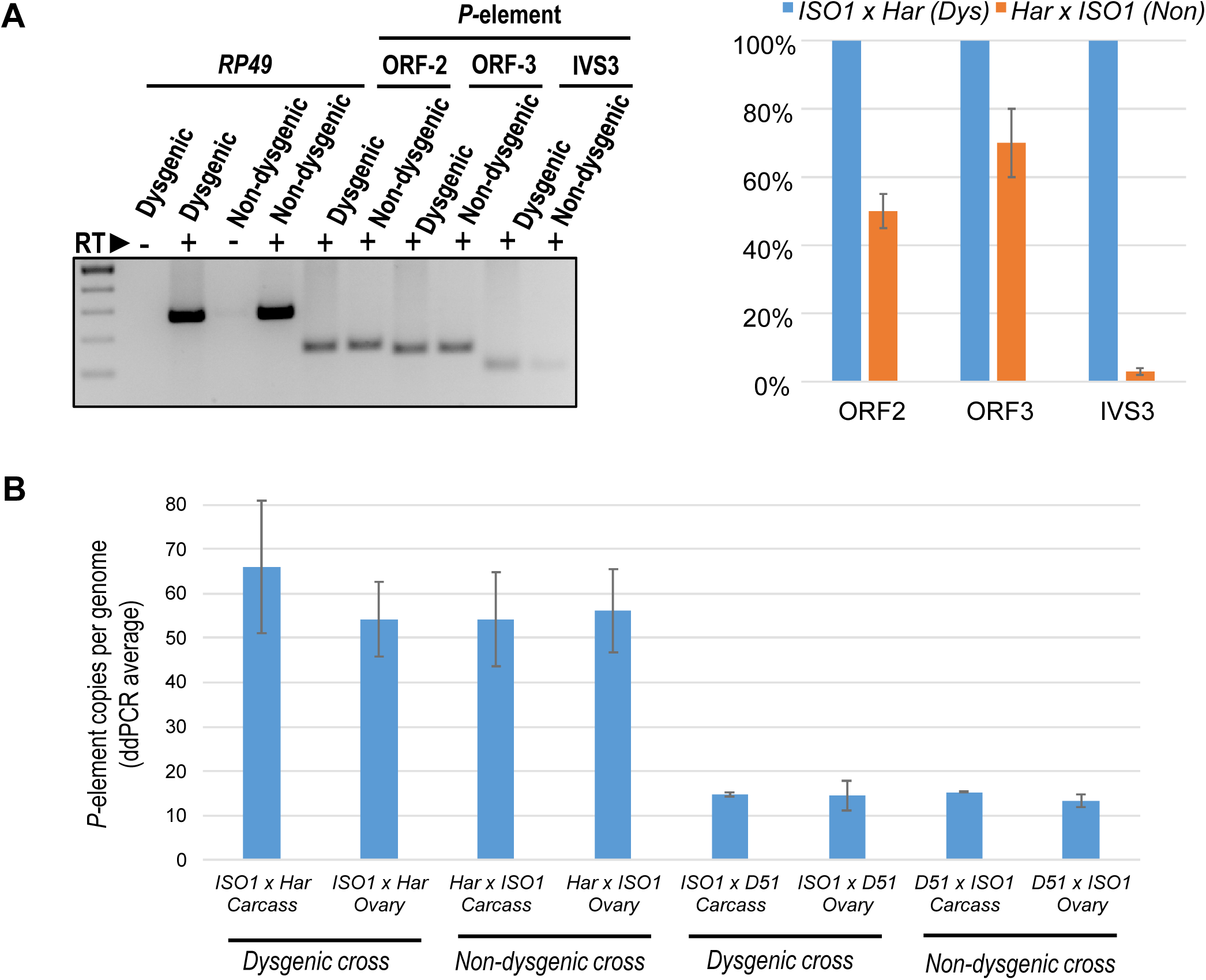
*P*-transposase RNA expression and *P*-copy number load is largely similar between Dysgenic and Non-dysgenic cross daughters. (A) *P-*element RNA expression in 3-5 days old F1 ovaries, *ISO1* x *Har* (Dysgenic) versus *Har x ISO1* (Non-dysgenic). Primers are specific to the ORF2 and ORF3 regions of the *P*-element, while the primers for IVS3 measure only the spliced product of IVS3 intron removal from the *P*-element mRNA. Relative quantifications of qPCR measurements was calculated from ΔΔCt with *RP49* as reference gene. (B) Digital droplet PCR quantitation of genomic *P*-element loads in daughter ovaries and carcasses from Dysgenic versus Non-dysgenic crosses. Error bars are standard deviation of 3 biological replicates of crosses.

**Figure 7-S1.**
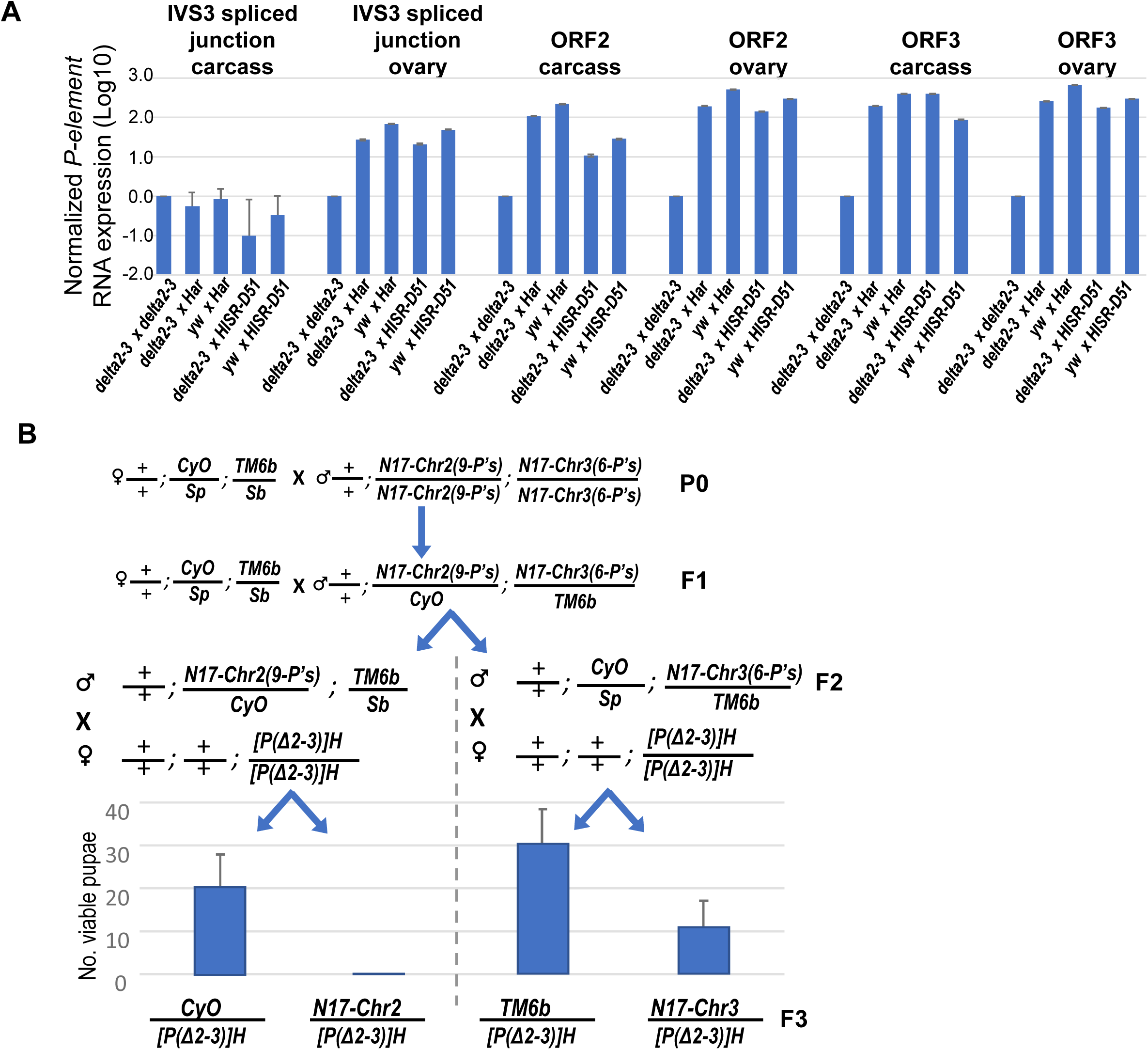
**Further examinations of *delta2-3 P*-transposase RNA expression and *delta2-3 P*-transposase capacity to cause pupal lethality with lower copy numbers of *Har-P’s*.** (A) The *P*-transposase mRNA expression is not silenced in the progeny of crosses between mothers expressing *delta2-3 P*-transposase (and *yw* control) with fathers that came from mothers expressing *P*-element-directed piRNAs. Relative quantifications of qPCR measurements was calculated from ΔΔCt with *RP49* as reference gene. (B) A genetic cross to assay how *Har-P* copy number and composition impacts the strong pupal lethality phenotype in a cross with the *delta2-3* P-transposase. The *HISR-N17* strain was first crossed to the double-balancer line *+/+; CyO/Sp; TM6b/Sb* to generate to males selected for *CyO and TM6b* that were then crossed again to the double-balancer to select F2 males with *Sb* or *Sp* traits. These F2 males were then each crossed to *[P(Δ2-3)]H* to generate viable pupae which emerged as F3 adult flies and were assessed for *CyO* and *TM6b*. The average and standard deviation of three biological replicates are plotted.

Tables S1 to S2.

## REFERENCES

1. Adams, M.D., Celniker, S.E., Holt, R.A., Evans, C.A., Gocayne, J.D., Amanatides, P.G., Scherer, S.E., Li, P.W., Hoskins, R.A., Galle, R.F., et al. (2000). The genome sequence of Drosophila melanogaster. Science 287, 2185–2195.

2. Anxolabehere, D., Nouaud, D., Periquet, G., and Tchen, P. (1985). P-element distribution in Eurasian populations of Drosophila melanogaster: A genetic and molecular analysis. Proceedings of the National Academy of Sciences of the United States of America 82, 5418–5422.

3. Aravin, A.A., Sachidanandam, R., Bourc’his, D., Schaefer, C., Pezic, D., Toth, K.F., Bestor, T., and Hannon, G.J. (2008). A piRNA pathway primed by individual transposons is linked to de novo DNA methylation in mice. Molecular cell 31, 785–799.

4. Bergman, C.M., Han, S., Nelson, M.G., Bondarenko, V., and Kozeretska, I. (2017). Genomic analysis of P elements in natural populations of Drosophila melanogaster. PeerJ 5, e3824.

5. Biemont, C., Ronsseray, S., Anxolabehere, D., Izaabel, H., and Gautier, C. (1990). Localization of P elements, copy number regulation, and cytotype determination in Drosophila melanogaster. Genet Res 56, 3–14.

6. Bingham, P.M., Kidwell, M.G., and Rubin, G.M. (1982). The molecular basis of P-M hybrid dysgenesis: the role of the P element, a P-strain-specific transposon family. Cell 29, 995–1004.

7. Black, D.M., Jackson, M.S., Kidwell, M.G., and Dover, G.A. (1987). KP elements repress P-induced hybrid dysgenesis in Drosophila melanogaster. The EMBO journal 6, 4125–4135.

8. Boussy, I.A., Healy, M.J., Oakeshott, J.G., and Kidwell, M.G. (1988). Molecular analysis of the P-M gonadal dysgenesis cline in eastern Australian Drosophila melanogaster. Genetics 119, 889–902.

9. Brennecke, J., Aravin, A.A., Stark, A., Dus, M., Kellis, M., Sachidanandam, R., and Hannon, G.J. (2007). Discrete small RNA-generating loci as master regulators of transposon activity in Drosophila. Cell 128, 1089–1103.

10. Brennecke, J., Malone, C.D., Aravin, A.A., Sachidanandam, R., Stark, A., and Hannon, G.J. (2008). An epigenetic role for maternally inherited piRNAs in transposon silencing. Science 322, 1387–1392.

11. Chung, W.J., Okamura, K., Martin, R., and Lai, E.C. (2008). Endogenous RNA interference provides a somatic defense against Drosophila transposons. Curr Biol 18, 795–802.

12. Clark, J.P., Rahman, R., Yang, N., Yang, L.H., and Lau, N.C. (2017). Drosophila PAF1 Modulates PIWI/piRNA Silencing Capacity. Curr Biol 27, 2718–2726 e2714.

13. Danecek, P., Auton, A., Abecasis, G., Albers, C.A., Banks, E., DePristo, M.A., Handsaker, R.E., Lunter, G., Marth, G.T., Sherry, S.T., et al. (2011). The variant call format and VCFtools. Bioinformatics 27, 2156–2158.

14. DePristo, M.A., Banks, E., Poplin, R., Garimella, K.V., Maguire, J.R., Hartl, C., Philippakis, A.A., del Angel, G., Rivas, M.A., Hanna, M., et al. (2011). A framework for variation discovery and genotyping using next-generation DNA sequencing data. Nature genetics 43, 491–498.

15. Engels, W.R., Benz, W.K., Preston, C.R., Graham, P.L., Phillis, R.W., and Robertson, H.M. (1987). Somatic effects of P element activity in Drosophila melanogaster: pupal lethality. Genetics 117, 745–757.

16. Engels, W.R., and Preston, C.R. (1979). Hybrid dysgenesis in Drosophila melanogaster: the biology of female and male sterility. Genetics 92, 161–174.

17. Erwin, A.A., Galdos, M.A., Wickersheim, M.L., Harrison, C.C., Marr, K.D., Colicchio, J.M., and Blumenstiel, J.P. (2015). piRNAs Are Associated with Diverse Transgenerational Effects on Gene and Transposon Expression in a Hybrid Dysgenic Syndrome of D. virilis. PLoS genetics 11, e1005332.

18. Ghildiyal, M., Seitz, H., Horwich, M.D., Li, C., Du, T., Lee, S., Xu, J., Kittler, E.L., Zapp, M.L., Weng, Z., et al. (2008). Endogenous siRNAs derived from transposons and mRNAs in Drosophila somatic cells. Science 320, 1077–1081.

19. Gloor, G.B., Preston, C.R., Johnson-Schlitz, D.M., Nassif, N.A., Phillis, R.W., Benz, W.K., Robertson, H.M., and Engels, W.R. (1993). Type I repressors of P element mobility. Genetics 135, 81–95.

20. Han, B.W., Wang, W., Zamore, P.D., and Weng, Z. (2015). piPipes: a set of pipelines for piRNA and transposon analysis via small RNA-seq, RNA-seq, degradome- and CAGE-seq, ChIP-seq and genomic DNA sequencing. Bioinformatics 31, 593–595.

21. Hancks, D.C., and Kazazian, H.H., Jr. (2012). Active human retrotransposons: variation and disease. Current opinion in genetics & development 22, 191–203.

22. Jackson, M.S., Black, D.M., and Dover, G.A. (1988). Amplification of KP elements associated with the repression of hybrid dysgenesis in Drosophila melanogaster. Genetics 120, 1003–1013.

23. Josse, T., Teysset, L., Todeschini, A.L., Sidor, C.M., Anxolabehere, D., and Ronsseray, S. (2007). Telomeric trans-silencing: an epigenetic repression combining RNA silencing and heterochromatin formation. PLoS genetics 3, 1633–1643.

24. Kawamura, Y., Saito, K., Kin, T., Ono, Y., Asai, K., Sunohara, T., Okada, T.N., Siomi, M.C., and Siomi, H. (2008). Drosophila endogenous small RNAs bind to Argonaute 2 in somatic cells. Nature 453, 793–797.

25. Kelleher, E.S. (2016). Reexamining the P-Element Invasion of Drosophila melanogaster Through the Lens of piRNA Silencing. Genetics 203, 1513–1531.

26. Khurana, J.S., Wang, J., Xu, J., Koppetsch, B.S., Thomson, T.C., Nowosielska, A., Li, C., Zamore, P.D., Weng, Z., and Theurkauf, W.E. (2011). Adaptation to P element transposon invasion in Drosophila melanogaster. Cell 147, 1551–1563.

27. Kidwell, M.G., Kidwell, J.F., and Sved, J.A. (1977). Hybrid Dysgenesis in DROSOPHILA MELANOGASTER: A Syndrome of Aberrant Traits Including Mutation, Sterility and Male Recombination. Genetics 86, 813–833.

28. Kidwell, M.G., Novy, J.B., and Feeley, S.M. (1981). Rapid unidirectional change of hybrid dysgenesis potential in Drosophila. J Hered 72, 32–38.

29. Kozeretska, I.A., Shulha, V.I., Serga, S.V., Rozhok, A.I., Protsenko, O.V., and Lau, N.C. (2018). A rapid change in P-element-induced hybrid dysgenesis status in Ukrainian populations of Drosophila melanogaster. Biol Lett 14.

30. Lau, N.C., Robine, N., Martin, R., Chung, W.J., Niki, Y., Berezikov, E., and Lai, E.C. (2009). Abundant primary piRNAs, endo-siRNAs, and microRNAs in a Drosophila ovary cell line. Genome research 19, 1776–1785.

31. Le Thomas, A., Stuwe, E., Li, S., Du, J., Marinov, G., Rozhkov, N., Chen, Y.C., Luo, Y., Sachidanandam, R., Toth, K.F., et al. (2014). Transgenerationally inherited piRNAs trigger piRNA biogenesis by changing the chromatin of piRNA clusters and inducing precursor processing. Genes & development 28, 1667–1680.

32. Li, H., and Durbin, R. (2010). Fast and accurate long-read alignment with Burrows-Wheeler transform. Bioinformatics 26, 589–595.

33. Marin, L., Lehmann, M., Nouaud, D., Izaabel, H., Anxolabehere, D., and Ronsseray, S. (2000). P-Element repression in Drosophila melanogaster by a naturally occurring defective telomeric P copy. Genetics 155, 1841–1854.

34. McKenna, A., Hanna, M., Banks, E., Sivachenko, A., Cibulskis, K., Kernytsky, A., Garimella, K., Altshuler, D., Gabriel, S., Daly, M., et al. (2010). The Genome Analysis Toolkit: a MapReduce framework for analyzing next-generation DNA sequencing data. Genome research 20, 1297–1303.

35. Moon, S., Cassani, M., Lin, Y.A., Wang, L., Dou, K., and Zhang, Z.Z. (2018). A Robust Transposon-Endogenizing Response from Germline Stem Cells. Developmental cell 47, 660–671 e663.

36. Mullins, M.C., Rio, D.C., and Rubin, G.M. (1989). cis-acting DNA sequence requirements for P-element transposition. Genes & development 3, 729–738.

37. O’Hare, K., and Rubin, G.M. (1983). Structures of P transposable elements and their sites of insertion and excision in the Drosophila melanogaster genome. Cell 34, 25–35.

38. Rahman, R., Chirn, G.W., Kanodia, A., Sytnikova, Y.A., Brembs, B., Bergman, C.M., and Lau, N.C. (2015). Unique transposon landscapes are pervasive across Drosophila melanogaster genomes. Nucleic acids research 43, 10655–10672.

39. Rasmusson, K.E., Raymond, J.D., and Simmons, M.J. (1993). Repression of hybrid dysgenesis in Drosophila melanogaster by individual naturally occurring P elements. Genetics 133, 605–622.

40. Rathke, C., Baarends, W.M., Jayaramaiah-Raja, S., Bartkuhn, M., Renkawitz, R., and Renkawitz-Pohl, R. (2007). Transition from a nucleosome-based to a protamine-based chromatin configuration during spermiogenesis in Drosophila. J Cell Sci 120, 1689–1700.

41. Robertson, H.M., and Engels, W.R. (1989). Modified P elements that mimic the P cytotype in Drosophila melanogaster. Genetics 123, 815–824.

42. Robertson, H.M., Preston, C.R., Phillis, R.W., Johnson-Schlitz, D.M., Benz, W.K., and Engels, W.R. (1988). A stable genomic source of P element transposase in Drosophila melanogaster. Genetics 118, 461–470.

43. Ronsseray, S., Lehmann, M., and Anxolabehere, D. (1989). Copy number and distribution of P and I mobile elements in Drosophila melanogaster populations. Chromosoma 98, 207–214.

44. Rubin, G.M., Kidwell, M.G., and Bingham, P.M. (1982). The molecular basis of P-M hybrid dysgenesis: the nature of induced mutations. Cell 29, 987–994.

45. Siebel, C.W., Kanaar, R., and Rio, D.C. (1994). Regulation of tissue-specific P-element pre-mRNA splicing requires the RNA-binding protein PSI. Genes & development 8, 1713–1725.

46. Simmons, M.J., Haley, K.J., Grimes, C.D., Raymond, J.D., and Fong, J.C. (2002a). Regulation of P-element transposase activity in Drosophila melanogaster by hobo transgenes that contain KP elements. Genetics 161, 205–215.

47. Simmons, M.J., Haley, K.J., Grimes, C.D., Raymond, J.D., and Niemi, J.B. (2002b). A hobo transgene that encodes the P-element transposase in Drosophila melanogaster: autoregulation and cytotype control of transposase activity. Genetics 161, 195–204.

48. Simmons, M.J., Raymond, J.D., Boedigheimer, M.J., and Zunt, J.R. (1987). The influence of nonautonomous P elements on hybrid dysgenesis in Drosophila melanogaster. Genetics 117, 671–685.

49. Simmons, M.J., Raymond, J.D., Rasmusson, K.E., Miller, L.M., McLarnon, C.F., and Zunt, J.R. (1990). Repression of P element-mediated hybrid dysgenesis in Drosophila melanogaster. Genetics 124, 663–676.

50. Simmons, M.J., Ryzek, D.F., Lamour, C., Goodman, J.W., Kummer, N.E., and Merriman, P.J. (2007). Cytotype regulation by telomeric P elements in Drosophila melanogaster: evidence for involvement of an RNA interference gene. Genetics 176, 1945–1955.

51. Srivastav, S.P., and Kelleher, E.S. (2017). Paternal Induction of Hybrid Dysgenesis in Drosophila melanogaster Is Weakly Correlated with Both P-Element and hobo Element Dosage. G3 (Bethesda) 7, 1487-1497.

52. Sytnikova, Y.A., Rahman, R., Chirn, G.W., Clark, J.P., and Lau, N.C. (2014). Transposable element dynamics and PIWI regulation impacts lncRNA and gene expression diversity in Drosophila ovarian cell cultures. Genome research 24, 1977–1990.

53. Tang, M., Cecconi, C., Bustamante, C., and Rio, D.C. (2007). Analysis of P element transposase protein-DNA interactions during the early stages of transposition. J Biol Chem 282, 29002–29012.

54. Teixeira, F.K., Okuniewska, M., Malone, C.D., Coux, R.X., Rio, D.C., and Lehmann, R. (2017). piRNA-mediated regulation of transposon alternative splicing in the soma and germ line. Nature 552, 268–272.

55. Wakisaka, K.T., Ichiyanagi, K., Ohno, S., and Itoh, M. (2017). Diversity of P-element piRNA production among M’ and Q strains and its association with P-M hybrid dysgenesis in Drosophila melanogaster. Mob DNA 8, 13.

56. Wei, G., Oliver, B., and Mahowald, A.P. (1991). Gonadal dysgenesis reveals sexual dimorphism in the embryonic germline of Drosophila. Genetics 129, 203–210.

57. Yamaguchi, K., Hada, M., Fukuda, Y., Inoue, E., Makino, Y., Katou, Y., Shirahige, K., and Okada, Y. (2018). Re-evaluating the Localization of Sperm-Retained Histones Revealed the Modification-Dependent Accumulation in Specific Genome Regions. Cell reports 23, 3920–3932.

58. Yang, G., Nagel, D.H., Feschotte, C., Hancock, C.N., and Wessler, S.R. (2009). Tuned for transposition: molecular determinants underlying the hyperactivity of a Stowaway MITE. Science 325, 1391–1394.

59. Yoshitake, Y., Inomata, N., Sano, M., Kato, Y., and Itoh, M. (2018). The P element invaded rapidly and caused hybrid dysgenesis in natural populations of Drosophila simulans in Japan. Ecol Evol 8, 9590–9599.

